# Welcoming back my arm: Affective touch increases body ownership following right hemisphere stroke

**DOI:** 10.1101/851055

**Authors:** Paul M. Jenkinson, Cristina Papadaki, Sahba Besharati, Valentina Moro, Valeria Gobbetto, Laura Crucianelli, Louise P. Kirsch, Renato Avesani, Nick S. Ward, Aikaterini Fotopoulou

## Abstract

Right hemisphere stroke can impair the ability to recognise one’s contralesional body parts as belonging to one’s self. The study of this so-called ‘disturbed sense of limb ownership’ (DSO) can provide unique insights into the neurocognitive mechanisms of body ownership. Here, we address a hypothesis built upon experimental studies on body ownership in healthy volunteers. These studies have shown that affective (pleasant) touch, an interoceptive modality associated with unmyelinated, slow-conducting C tactile afferents, has a unique role in the sense of body ownership. Here we systematically investigated whether affective touch stimulation could increase body ownership in patients with DSO following right hemisphere stroke. An initial feasibility study in 16 adult, acute stroke patients enabled us to optimise and calibrate an affective touch protocol to be administered by the bedside. The main experiment, conducted with a different sample of 26 right hemisphere patients, assessed changes in limb ownership elicited following self-(patient) versus other-(experimenter) generated tactile stimulation, using a velocity known to optimally activate C-tactile fibres (i.e. 3cm/s), and a second velocity that is suboptimal for C-tactile activation (i.e. 18cm/s). We further examined the specificity and mechanism of observed changes in limb ownership in secondary analyses looking at (1) the influence of perceived intensity and pleasantness of touch, (2) touch laterality, and (3) level of DSO on ownership change, as well as (4) changes in unilateral neglect arising from touch. Findings indicated a significant increase in limb ownership following experimenter-administered, CT-optimal touch. Voxel-based Lesion-Symptom Mapping (VLSM) identified damage to the right insula and, more substantially, the right corpus callosum, associated with a failure to increase body ownership following experimenter-administered, affective touch. Our findings suggest that affective touch can increase the sense of body-part ownership following right hemisphere stroke, potentially due to its unique role in the multisensory integration processes that underlie the sense of body ownership.

## 1. Introduction

Several disturbances of body awareness occur after right-hemisphere stroke, including asomatognosia (i.e. feelings of non-belonging or non-recognition of the limb) and somatoparaphrenia (delusional ideas of disownership; see Jenkinson *et al*., 2018). These disturbances reveal that our seemingly effortless sense of body ownership (i.e. the sense that *my* body belongs to *me*) is actually supported by distinct neurocognitive mechanisms that warrant scientific study. In addition, these symptoms present a significant but unmet clinical challenge, with body unawareness being associated with poor engagement with rehabilitation, longer hospitalisation and poorer prognosis (see Jenkinson *et al*., 2011, Besharati *et al*., 2014*a*, *b*).

Existing research into the mechanisms of body ownership in healthy individuals has identified dynamic integration and weighting processes involving several exteroceptive (e.g. vision) and interoceptive signals (i.e. representing the physiological state of the body; see Craig, 2003). Amongst these signals, a specific type of pleasant touch (hereafter referred to as affective touch), plays a significant role (Crucianelli *et al*., 2013, 2017; Lloyd *et al*., 2013; van Stralen *et al*., 2014; Ponzo *et al*., 2018). Affective touch involves slow conducting, unmyelinated C-tactile afferent nerve fibres (CT afferents), which are thought to project mainly to the posterior insula via a distinct ascending pathway (Olausson *et al*., 2002; Mcglone *et al*., 2012), associated mainly with interoceptive rather than exteroceptive modalities (Craig, 2009; Ceunen *et al*., 2016). Microneurography studies have shown that CT-afferents, located on the hairy skin of the body, respond preferentially to slow (i.e. velocities between 1 and 10cm/s), dynamic, light-pressure, touch (Vallbo and Hagbarth, 1968; Nordin, 1990; Vallbo *et al*., 1993, 1999). Moreover, their activation is positively correlated with subjective ratings of tactile pleasure in healthy subjects (Löken *et al*., 2009). Importantly, work on peripheral neuropathies, as well as functional neuroimaging studies in healthy participants (see Morrison, 2016 for a review and meta-analysis) supports distinct functional roles between large diameter, fast-conducting, myelinated afferent fibres (Aβ-fibres) and the CT-afferent system. For example, patients with a genetic mutation that reduces CT-afferent density reported reduced pleasantness in response to CT-optimal touch, while discriminatory aspects of touch remained intact (Morrison *et al*., 2011). By contrast, a patient with selective loss of large, myelinated afferents showed the reverse pattern, with a faint but preserved sensation of pleasant touch in response to light, CT-optimal stoking touch on the forearm (Olausson *et al*., 2002).

Our group was the first to show that applying CT-optimal, affective touch during multisensory illusions of body-part ownership, such as the classic rubber hand illusion (Botvinick and Cohen, 1998), enhances ownership (Crucianelli *et al*., 2013). Several other studies and groups have now replicated this finding (Lloyd *et al*., 2013; van Stralen *et al*., 2014; Crucianelli *et al*., 2017; Panagiotopoulou *et al*., 2017; Ponzo *et al*., 2018) and further investigated different explanations of this effect. First, it appears that affective congruency between different modalities (e.g. seeing a soft, pleasant material and feeling one via touch) increases feelings of ownership during multisensory integration, over and above other amodal properties such as temporal and spatial congruency (Filippetti *et al*., 2019). Moreover, CT-optimal, affective touch can reduce sensations of ‘deafference’ during asynchronous modulation (Panagiotopoulou *et al*., 2017). ‘Deafference’ in this context refers to the unpleasant and numb feelings about one’s own body caused by the temporal mismatch (a cross-modal property the brain uses to bind sensations together during multisensory integration) between seen and felt tactile stimulation (Longo *et al*., 2008; Lesur *et al*., 2019). Thus, it is possible that CT-optimal, affective touch may also enhance body ownership of the affected arm in right-hemisphere patients with a disturbed sense of limb ownership (DSO; Baier and Karnath, 2008; Jenkinson *et al*., 2018). Although DSO and impaired somatosensation are known to dissociate (Moro *et al*., 2004; Vallar and Ronchi, 2009), to our knowledge these studies have only assessed discriminatory and spatial aspects of tactile perception. Thus, no study to date has examined whether spontaneous feelings of ‘deafference’, such as the sensation that the arm is abnormally cold, numb or painful, may contribute to DSO.

Although it can be hard to systematically sample such subjective, spontaneous feelings by the bedside, in a previous study we showed that DSO may be at least partly explained as sensations about the affected arm, which cannot be explained by existing top-down expectations of selfhood (Martinaud *et al*., 2017). Specifically, we found that almost all of our sample of patients with right perisylvian lesions (N = 31), with and without DSO symptoms, experienced feelings of ownership over a rubber hand within 15 seconds and without any tactile stimulation (a phenomenon termed visual capture of ownership, VOC). Paradoxically, the subset of these patients that had clinical somatoparaphrenia denied the ownership of their own arm, even when they accepted (mistakenly) the ownership of a rubber hand that was placed at the same position on the left hemispace. Thus, DSO is not merely a matter of being unable to attribute a seen left arm positioned in the left hemispace to the self, but rather it is possible that some unexpected sensation or feelings about the arm leads patients to infer that this arm cannot possibly be their own (for the wider theoretical context of this hypothesis see (Fotopoulou, 2015; Crucianelli *et al*., 2019). Unfortunately, in practice it can be difficult to reliably assess the presence of such spontaneous deafference sensations and their precise role in arm ownership. However, a possible alternative way to test these ideas is to experimentally *reduce* such sensations using affective touch stimulation (which as aforementioned can attenuate experimentally-induced feelings of deafference; Panagiotopoulou *et al*., 2017), and observe the effect on body-part ownership. Moreover, a previous single case study reported that gentle touch (not controlled for CT-fibre activation) increased feelings of arm belonging and related emotional attitudes in a right-hemisphere stroke patient with DSO (van Stralen *et al*., 2011).

A second aspect of touch, which may moderate its effect on body ownership and also requires examination, is whether touch is self-generated or originates from another person (externally-generated touch). Traditionally, self-generated touch has been associated with attenuation of the resulting tactile stimulation, leading to the well-known phenomenon that we cannot tickle ourselves (Blakemore *et al*., 1998*a*). Recent work in healthy subjects further shows how body ownership influences this sensory attenuation during the rubber hand illusion (Kilteni and Ehrsson, 2017). However, other research highlights the enhancing effect of self-touch on sensory perception (see Valentini *et al*., 2008; Ackerley *et al*., 2012), including in patients with right hemisphere stroke (White *et al*., 2010) and those with disturbed body ownership (see van Stralen *et al*., 2011). Attentional modulation and temporal expectation are proposed mechanisms for the observed self-enhancement effect. However, while the perceptual effects and neural basis of self-versus other-generated, CT-optimal, affective touch have been explored by our group (Gentsch *et al*., 2015) and others (e.g. Boehme *et al*., 2019), to our knowledge no group study has assessed the potentially beneficial effects that affective, self-versus other-generated touch may have on body ownership in stroke patients.

On the basis of the above, we aimed to examine how affective touch influences body-part ownership in patients with DSO. We predicted that caress-like touch delivered at CT-optimal velocities, would enhance limb ownership in right hemisphere stroke patients with DSO. We also included in our study a CT-suboptimal, fast-touch control condition in order to establish whether the mechanism underlying any touch-based increase in limb ownership is mediated by the CT system, as opposed to being the result of more generic touch-based, or attentional effects. This faster, but otherwise identical, touch is known to not activate the CT system optimally and is typically judged as more intense but less pleasant than touch activating the CT system (see e.g. Case *et al*., 2016). We also examined how touch delivered by the patient themselves (self-touch) versus another person (other-touch) might moderate these effects. Because of the opposing effects reported in the extant literature, we did not make a-priori predictions regarding which kind of touch (self or other) would produce greater changes in body-part ownership. In secondary, exploratory manipulations and statistical analyses we aimed to examine the specificity of any touch effects we observed, by investigating the relationship between CT-touch and CT-suboptimal touch perception (intensity and pleasantness ratings), levels of DSO and changes in ownership. We also implemented two control manipulations to examine the specificity of CT-touch effects. First, we examined the potential effect of affective touch on another common symptom following right hemisphere stroke, namely unilateral neglect. Second, we assessed whether changes in arm ownership result from a general, positive valence effect, by applying affective touch to the non-affected, ipsilateral (right) arm and measuring changes in ownership of the contralesional (left) arm. Finally, in exploratory lesion analyses we examined the underlying neuroanatomy of DSO and changes in DSO consequent to touch, which have not been examined in previous studies.

## 2. Materials and method

### 2.1. Patients

Forty-two, acute, right-hemisphere stroke patients were recruited by screening consecutive admissions at stroke-rehabilitation wards located in London, UK and Verona, Italy as part of larger, ongoing projects on body awareness following right-hemisphere stroke. Inclusion criteria were: (i) right-hemisphere lesion confirmed by clinical neuroimaging (CT or MRI), (ii) dense contralesional hemiplegia (i.e. power < 1; MRC scale; see Assessments below), and (iii) < 4 months from symptom onset, (iv) right handed. Exclusion criteria comprised: (i) previous history of neurological or psychiatric illness, ii) < 7 years of education, (iii) medication with significant cognitive or mood side-effects, and (iv) language impairments that precluded completion of assessments. Sixteen of these patients were recruited in the UK to an extensive feasibility study, which we conducted prior to the main experiment. In this initial feasibility study, we explored whether patients were able to tolerate a relatively long, touch protocol, perceive the difference between affective and neutral touch, and show some difference in their body representation as a result of touch. This study informed our predictions and, importantly, allowed us to develop a robust but also clinically feasible experimental protocol for the main study (full details reported in Supplementary Materials). The remaining 28 patients were recruited consecutively from both the UK (n=21) and Italy (n=7) and took part in only the main experiment. We subsequently excluded 2 patients from the final data analysis: one due to an experimenter error leading to incorrect administration of the experimental protocol, and another containing a large amount of missing data (i.e. completed only 1/4 experimental conditions) due to patient fatigue causing termination of the test protocol; therefore, a total of 26 patients contributing to the final, main analysis (see Table 1 for full details). The study was approved by the relevant University and National Health Service (UK) ethical committees, and patient consent obtained according to the Declaration of Helsinki.

**Table 1.** Clinical and neuropsychological characteristics of patients.

|  | *Experimental Baseline DSO Score (Range:0-3)* |  |  |  |  |
| --- | --- | --- | --- | --- | --- |
|  | 0<br>(N = 6 ) | 0.5<br>(N = 5) | 1<br>(N = 5) | 2<br>(N =10 ) | *K-W test* |
| Males:Females | 1:4<sub>a</sub> | 2:2<sub>a</sub> | 2:2<sub>a</sub> | 7:3 |  |
| Age (years) | 76<sub>a</sub><br>(55.50 – 78.50) | 59.50<sub>a</sub><br>(41.25 – 77) | 54.50<sub>a</sub><br>(39 – 73.75) | 66<sub>a</sub><br>(58.75 – 73.50) | H(3)=1.65,<br>p = .649 |
| Education (years) | 12<sub>a</sub><br>(8.5 – 14) | 15<sub>a</sub><br>(12.75 – 15) | 9<sub>c</sub><br>(5 – 13) | 9<sub>a</sub><br>(6.5 – 13.5) | H(3) = 4.71,<br>p = .195 |
| Days from stroke onset | 12<sub>a</sub><br>(2 – 40) | 16<sub>a</sub><br>(6.75 – 26) | 19<sub>a</sub><br>(15.5 – 77.25) | 14.5<br>(4.25 – 44.5) | H(3) = 1.17,<br>p = .759 |
| MRC Left upper limb | 0<sub>a</sub><br>(0 – 0.5) | 0<sub>b</sub><br>(0 – 0) | 0<sub>a</sub><br>(0 – 0) | 0<br>(0 – 0.25) | H(3) = 1.55,<br>p = .671 |
| Orientation (person, time & place) (max 3) | 3<sub>a</sub><br>(3 – 3) | 3<sub>b</sub><br>(2 – 3) | 2.5<sub>b</sub><br>(2 – 3) | 2.5<sub>b</sub><br>(2 – 3) | H(3) = 4.01,<br>p = .261 |
| Digit span forwards (max number repeated) | 7<sub>b</sub><br>(6.25 – 7.75) | 6.5<sub>a</sub><br>(5.25 – 7.75) | 6<sub>a</sub><br>(5.25 – 6.75) | 6<sub>a</sub><br>(4.5 – 7) | H(3) = 2.59,<br>p = .460 |
| Digit span backwards (max number repeated) | 4.5<sub>b</sub><br>(2.5 – 7.25) | 4<sub>a</sub><br>(2.5 – 4.75) | 4<sub>a</sub><br>(2.5 – 4.75) | 5<sub>a</sub><br>(1.5 – 5.5) | H(3) = 0.76,<br>p = .88 |
| MoCA (max 30) | 21<sub>a</sub><br>(15 – 22.5) | 21.5<sub>a</sub><br>(19.25 – 24.5) | 19.5<sub>a</sub><br>(14.5 – 25.25) | 22<sub>b</sub><br>(12 – 24) | H(3) = 0.59,<br>p = .899 |
| MoCA memory (max 5) | 2<sub>c</sub><br>(0 – 5) | 4<sub>b</sub><br>(2 – 5) | 3<sub>c</sub><br>(2 – 4) | 3<sub>e</sub><br>(0 – 4) | H(3) = 1.24,<br>p = .743 |
| FAB total score (max 18) | 11<sub>b</sub><br>(4.75 – 12.75) | 11<sub>b</sub><br>(11 – 13) | 14<sub>b</sub><br>(11 – 15) | 7<sub>g</sub><br>(1 – 14) | H(3) = 3.04,<br>p = .386 |
| Premorbid IQ-WTAR (max 50) | 26<sub>e</sub> (26 – 26) | 36<sub>c</sub> (35 – 37) | - | - | H(1) = 1.5, p = .221 |
| Comb/razor test (% bias = left – right strokes/ left + ambiguous + right strokes) | -14.29<sub>a</sub><br>(-40.52 - -3.75) | -0.71<sub>a</sub><br>(-51.32 – 12.50) | -20.36<sub>a</sub><br>(-46.87 – 24.82) | -14.71<sub>a</sub><br>(-29.38 - -1.53) | H(3) = .90, p = .827 |
| Bisiach one item test (max 3) | 0<sub>a</sub><br>(0 – 1.5) | 0<sub>b</sub><br>(0 – 1) | 1<sub>b</sub><br>(0 – 1) | 0<sub>d</sub><br>(0 – 0.25) | H(3) = 2.04,<br>p = .564 |
| BIT - Line crossing (max 36) | 35<sub>a</sub><br>(12 – 36) | 26<sub>a</sub><br>(5.25 – 35.5) | 30<sub>b</sub><br>(22 – 36) | 17<sub>a</sub><br>(6 – 23) | H(3) = 3.71,<br>p = .295 |
| BIT - Star cancelation (max 54) | 19.5<sub>b</sub><br>(12 – 45.75) | 15<sub>b</sub><br>(6 – 54) | 27<sub>a</sub><br>(3.5 – 49.75) | 9.5<sub>d</sub><br>(5 – 33.75) | H(3) = 1.20,<br>p = .752 |
| BIT - Copy (max 3) | 0<sub>c</sub><br>(0 – 0) | 1.5<sub>c</sub><br>(0 – 3) | 2<sub>c</sub><br>(1 – 3) | 0<sub>f</sub><br>(0 – 2.25) | H(3) = 4.13,<br>p = .248 |
| BIT - Representational drawing (max 3) | 0<sub>b</sub><br>(0 – 1.5) | 1<sub>b</sub><br>(0 – 3) | 0<sub>c</sub><br>(0 – 0) | 0<sub>d</sub><br>(0 – 2.25) | H(3) = 2.23,<br>p = .527 |
| BIT - Line bisection (max 9) | 4<sub>a</sub><br>(1.5 – 9) | 3<sub>b</sub><br>(2 – 6) | 4<sub>b</sub><br>(0 – 4) | 0.5<sub>b</sub><br>(0 – 7.5) | H(3) = 1.36,<br>p = .715 |
| HADS depression (max 21) | 9<sub>c</sub><br>(2 – 14) | 6<sub>b</sub><br>(4 – 8) | 7<sub>c</sub><br>(3 – 11) | 3<sub>g</sub><br>(2 – 5) | H(3) = 2.10,<br>p = .552 |
| HADS anxiety (max 21) | 10<sub>c</sub><br>(8 – 17) | 5<sub>b</sub><br>(3 – 7) | 10.5<sub>c</sub><br>(10 – 11) | 7<sub>g</sub><br>(3 – 7) | H(3), = 7.79,<br>p = .051 |
| Nottingham (left arm: min 0 = no sensation, max 2 = no deficit) | 0<sub>a</sub><br>(0 – 1.5) | 0<sub>a</sub><br>(0 – 0) | 0.5<sub>a</sub><br>(0 – 1) | 0<sub>b</sub><br>(0 – 1.75) | H(3) = 2.27,<br>p = .519 |
| Bisiach motor awareness scale (max unawareness = 3) | 1.5<sub>a</sub><br>(0.5 – 3) | 0<sub>a</sub><br>(0 – 1.5) | 0.5<sub>a</sub><br>(0 – 2.5) | 0.75<br>(0 – 3) | H(3) = 2.16,<br>p = .539 |
| Feinberg awareness scale (max unawareness = 10) | 6.5<sub>a</sub><br>(2.75 – 10) | 1.75<sub>a</sub><br>(0.25 – 5.5) | 1.5<sub>a</sub><br>(0 – 6.38) | 2.5<sub>c</sub><br>(1 – 8) | H(3) = 3.44,<br>p = .329 |

*Note: Reported values are median and IQR (in parentheses; 25<sup>th</sup> – 75<sup>th</sup> quartile) given non-normal*
*distribution of data. Dashed lines indicate no data available. Values calculated with missing data: a = group n-1; b = group n-2; c = group n-3; d = group n-4; e = group n-5; f = group n-6; g = group n-7. See section 2.2 ‘clinical and cognitive assessment’ for details of measures reported in this table.*

### 2.2. Clinical and cognitive assessment

Patients underwent a neurological and neuropsychological examination comprising: a clinical assessment of left-right orientation (during which patients were asked to identify visually or use their unaffected side to point to/touch their own/the experimenter’s left/right hand, ear and leg), assessment of motor ability (power) of the left upper and lower limbs (Medical Research Council scale, MRC; Guarantors of Brain, 1986), upper and lower visual fields and tactile extinction on the upper and lower limbs (using the Bisiach *et al*., 1986 technique), light touch perception on the left arm (Revised Nottingham Sensory Assessment; rNSA; Lincoln *et al*., 1998), orientation in time and space as well as general cognitive ability (Montreal Cognitive Assessment; MoCA; Nasreddine *et al*., 2005), estimated premorbid intelligence (Weschler Test of Adult Reading; Wechsler, 2001), working memory (using the verbal digit span subtest of the Wechsler Adult Intelligence Scale III; Wechsler, 1997), visuospatial neglect (using five subtests [star cancellation, line bisection, line crossing, copy, and clock drawing] of the Behavioural Inattention Test; Halligan *et al*., 1991), personal neglect (one-item test; Bisiach *et al*., 1986; and comb/razor test; McIntosh *et al*., 2000), anosognosia for hemiplegia (Bisiach *et al*., 1986; Feinberg *et al*., 2000), executive function (Frontal Assessment Battery; FAB; Dubois *et al*., 2000), as well as brief assessment of depression and anxiety (Hospital Anxiety and Depression Scale; HADS; Zigmond and Snaith, 1983).

### 2.3. Experimental design and measures

To examine the effect of touch on limb disownership, we employed a 2 (touch valence: affective vs. neutral) x 2 (touch administrator: self vs. other) repeated-measures design. The valence of touch (affective vs. neutral) was manipulated using two different velocities, such that touch was administered using either a slow, CT-optimal (3cm/s) or fast, CT-suboptimal (18cm/s) touch, which are typically rated as relatively high vs. low in terms of tactile pleasantness and can be reliably distinguished from each other (see Crucianelli *et al*., 2013, 2017; Ponzo *et al*., 2018). This touch was administered by either the patient (self) or experimenter (other) in a block design (see Procedure), whereby four affective or neutral touch trials were delivered per block (plus one sham trial in which the movement was simulated but no touch was delivered to the forearm). The order of conditions was only partly counterbalanced, based on what our sample size and practical, bedside considerations allowed; thus we examined the possible influence of these order effects in specific analyses, below.

The primary dependent variable was “ownership change”. The level of disownership during the experimental session was assessed prior to any touch conditions (i.e. baseline disownership) and after each touch condition (i.e. post self-neutral, other-neutral, self-affective and other-affective touch) by asking three questions designed to assess arm recognition, feelings of belonging, and existence (as discussed by Jenkinson *et al*., 2018): (1) “Is this [pointing to the patient’s left arm] your own arm?”, (2) “Does it ever feel like this [patient’s left] arm does not belong to you/is not really yours?”, (3) “Do you ever feel that your left arm is missing/has disappeared?”. In order to avoid repetition and loss of attention, the exact wording used was varied on a trial-by-trial basis as indicated above (see Supplementary Materials for further details of the feasibility study used to develop this procedure). Patient responses to each question were recorded verbatim and scored by the experimenter using an adapted (i.e. to assess disownership rather than awareness) version of the established Feinberg *et al*., (2000) method, where higher scores indicated greater levels of disownership: 0 = no disownership; 0.5 = partial disownership; 1 = disownership. A second, independent researcher verified the scoring of all responses. In order to calculate “ownership change” we totalled the raw scores (total disownership score minimum = 0, maximum = 3), and then subtracted the post-touch disownership score from the pre-touch baseline (i.e. baseline score - post-touch score), so that a post-touch increase in ownership was indexed by a positive ownership change score, while any increase in disownership was indexed by a negative ownership change.

In order to explore the specificity and mechanism of the CT touch effect we also conducted a number of control tasks in a subset of patients and trials (constrained by fatigue and other practical considerations, these were merely exploratory investigations; see Supplementary Materials for details), to examine (1) the effect of touch on visuospatial and personal neglect and awareness of neglect, (2) whether touch applied to the *right* arm would influence disownership of the *left* limb, and the effect of CT and CT-suboptimal touch perception on ownership change.

### 2.4. Procedures

Touch was administered to the dorsal surface of the forearm (in a preselected area between the wrist and elbow crease; approximately 18cm), by a female experimenter using a soft make-up brush made from natural hair (Natural hair Blush Brush, No. 7, The Boots Company). Visual feedback of the touched arm was blocked either by asking the patient to wear a blindfold, or (if patients refused to wear a blindfold) by closing their eyes while an assistant experimenter also held a cardboard carton over the touched arm to obscure a direct view. Each touch *trial* comprised four strokes in alternating directions, from the elbow to the wrist and vice versa, with a pause of 1s between each stroke to reduce fibre fatigue. Slow touch strokes at a velocity of 3cm/s were administered in six seconds over the 18cm long area and fast strokes over the same area lasted 1 second. Each *block* comprised four trials of the same prespecified touch velocity (i.e. 4 x 4 touch trials = 16 strokes per block), and one randomly inserted sham trial, during which the brush was held several centimetres above the touch location and moved in order to mimic the slow/fast touch, but did not make contact with the patient’s skin. See Figure 1 for a schematic of the experiment.

**Figure 1.**
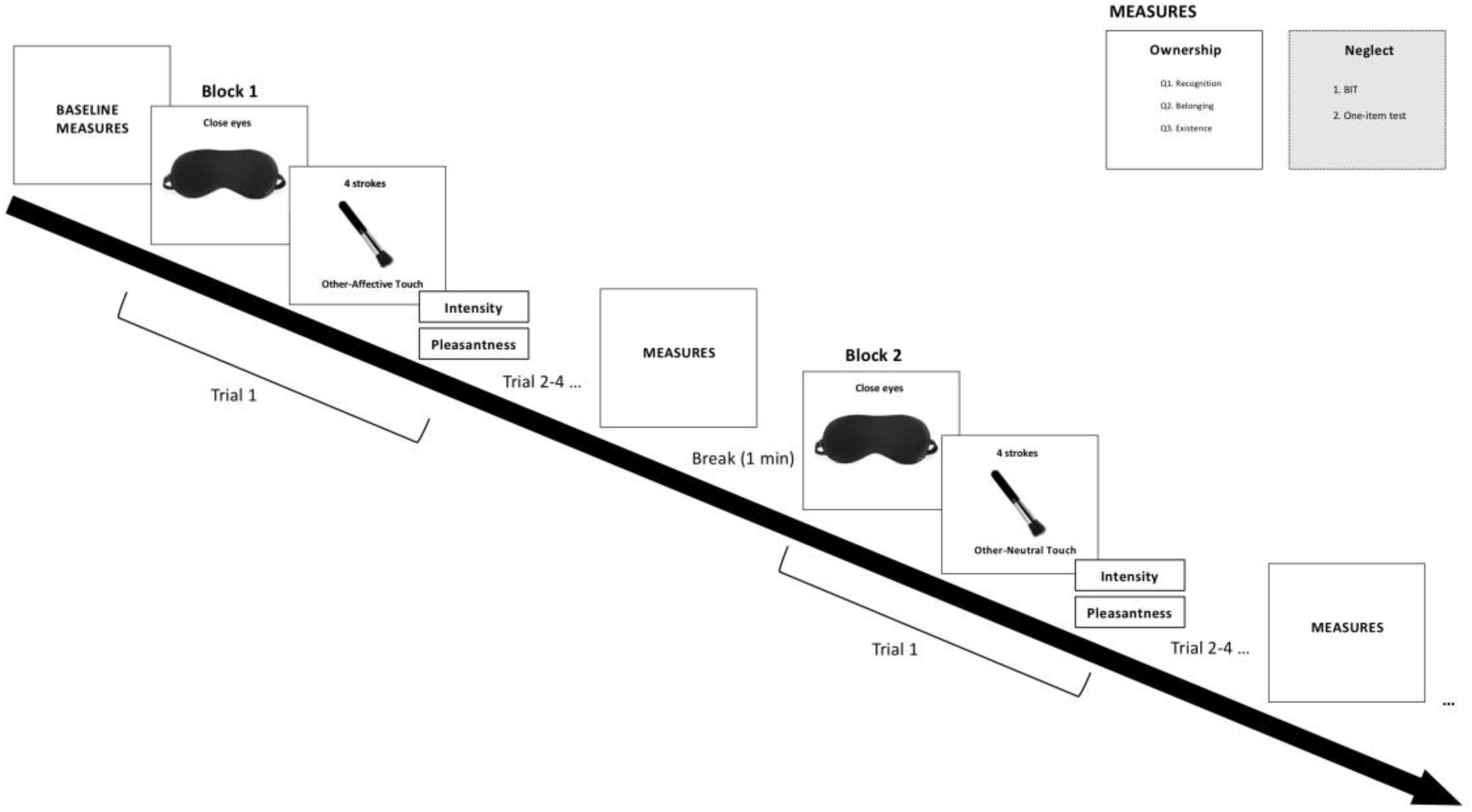
Schematic of the study procedure. Patients first completed a pre-touch baseline measure of ownership, followed by a block of touch (self-affective, other-affective, self-neutral or other-neutral), and a post-touch measure of arm ownership. A sham (no-touch) trial (not shown above for simplicity) was randomly inserted into each block. Neglect was assessed in a subsample of patients and only pre-post affective touch conditions. Four main blocks completed in total (self-affective, other-affective, self-neutral, other-neutral). Right arm control touch blocks were administered following the same method (without neglect assessment) in a subset of patients.

During a pre-experimental phase, patients were introduced to the stroking apparatus, procedure (i.e. slow, fast and sham trials) and vertical response scale (to minimize the effects of neglect; we also always ensured the participants could see the scale and read it aloud to facilitate them). Understanding of the response scale was also established by asking patients to rate the pleasantness of various physical and conceptual items (i.e. to be touched by a thorn / cotton on the skin, to win the lottery, to lose your keys, to be praised by someone you love), with any misunderstandings clarified prior to continuing to the main experimental protocol. One trial of slow/fast touch was also administered during this pre-experimental stage to establish if the patient could perceive touch on the left arm.

During *other*-*touch* blocks, the experimenter held the brush and touched the patient in the required location (left or right arm) using the appropriate velocity (slow or fast). By contrast, during *self-touch* blocks, the patient was physically guided (by the experimenter holding/supporting the patient’s hand/arm and brush when necessary) to touch the relevant location on the contralateral arm using the appropriate speed (for this method of self-touch, see White *et al*., 2010). This physical support was provided during all self-touch trials (i.e. both when the unaffected [right] and hemiplegic [left] arms were the ‘active’ arms delivering the touch) to ensure consistency between self and other stroking in terms of velocity and pressure of stroking. Immediately after each trial (4 strokes) the patient was asked to rate the touch intensity and pleasantness. Ownership was assessed prior to any touch (i.e. pre-touch baseline), and again following each touch block. Neglect was assessed before and after each block of affective touch only, due to time and fatigue constraints. A one-minute break was introduced between blocks, with longer breaks given where necessary (i.e. patient appeared fatigued or requested a break).

### 2.5. Statistical analyses

#### 2.5.1. Main Experimental Measure – Ownership Change

Our main analysis aimed to determine whether and to what extent different types of touch would increase body ownership; therefore, only patients with some baseline level of disownership on the day of the experimental session (i.e. baseline DSO+ patients; n = 20) were suitable for the main analysis. Patients not expressing any DSO during the baseline assessment on that day (DSO-) were not appropriate for this analysis since they were asymptomatic for the primary behaviour of interest (i.e. they did not show disownership during the session) and were thus excluded from it; however, these patients provided relevant comparison data for feasibility, comprehension, suggestibility and other practical considerations applying to our experimental procedures in an acute stroke setting, as well as clinical and neuropsychological measures, lesion analysis and experimental control conditions (described in Supplementary Materials).

We used both frequentist and Bayesian statistics to assess the observed effects, depending on the aim and hypothesis in each case. The complementary use of these two statistical approaches is recommended by a number of authors to facilitate a fuller understanding of the data (see e.g. Howard *et al*., 2000; Dienes, 2014; Dienes and Mclatchie, 2018; Quintana and Williams, 2018). For frequentist statistical inference, we assessed normality via visual inspection of histograms and the Shapiro-Wilks test. Two patients did not complete one of the four experimental conditions. We therefore analysed the experimental data using both pairwise (i.e. case-by-case) and listwise deletion methods for dealing with missing data, in order to assess the impact of this missing data on the experimental findings. Bayesian statistics were performed in order to allow further interpretation of the observed effects, in particular, the extent to which data provided support for the alternative versus null hypotheses. Bayes Factors (BF_10_) indicate the relative strength for the null versus alternative hypotheses (i.e. the number of times more likely the data are under the alternative than the null hypothesis), and were used as a means of interpreting evidence for each hypothesis, using benchmarks provided by Jeffreys (1961). We interpreted a BF_10_ > 3 as substantial evidence for the alternative hypothesis, a BF_10_ < 0.3 as substantial evidence in favour of the null hypothesis, and BF_10_ > 0.3 < 3.0 as insensitive, weak or anecdotal evidence for either hypothesis (see Dienes, 2014; Quintana and Williams, 2018).

We examined the extent to which touch modified body ownership using parameter estimates (95% confidence [frequentist] / credibility [Bayesian] intervals), and calculating non-parametric / Bayesian, one-sample t-tests (owing to the limited sample size and non-normal distribution; Shapiro-Wilks test), testing the null hypothesis of zero change in body ownership as a result of the touch applied (i.e. one sample test value = 0). We corrected for multiple comparisons using a Bonferroni-corrected alpha of 0.0125 (i.e. 0.05 / 4; applicable to frequentist statistics only). Although we expected touch to increase ownership in patients with DSO+, we could not rule out the possibility that touch might reduce body ownership, and so we performed all statistics using 2-tailed tests. All behavioural analyses were performed using JASP (JASP Team, 2019). Figures for behavioural data were generated in R (R Core Team, 2013) using ggplot 2 (Wickham, 2016) and in MRICron (Rorden and Brett, 2000; Rorden *et al*., 2007) for lesion analysis.

#### 2.5.2. Clinical and neuropsychological variables

Clinical and neuropsychological data were summarised by grouping patients according to their score on the experimental measure of DSO taken during the pre-touch baseline of the experimental session. As is typical in neuropsychological research with acute stroke patients, not all assessments were fully completed by all patients, owing to discharge, scheduling issues, or patients becoming too ill to complete assessments. In these cases, data are included where available and indicated where missing. Differences in the clinical and neuropsychological variables between these groups were analysed for exploration only, given that these groupings comprised small patient numbers and were not used as the basis for our main experimental manipulation. We used non-parametric tests (Kruskal-Wallis test), and an alpha of 0.01 to account for false positives arising from multiple comparisons, whilst also avoiding being overly conservative.

#### 2.5.3. Control Analyses

A number of control tasks were conducted in a subset of our patients due to practical constraints (as detailed above and in Supplementary Materials) to look at the possible influence of order, unilateral neglect, perceived pleasantness and intensity of CT and CT-suboptimal touch, and touch lateralisation on body ownership change. There were no order effects and no significant relationship between touch pleasantness or intensity and baseline levels of body ownership or body ownership change. Moreover, our touch manipulation had no effect on neglect, nor did a control application of the touch protocol to the right arm affect the ownership of the affected, left arm in our patients, confirming the specificity of our main findings. Thus, as none of these factors were found to relate to ownership changes in our study (our main measure), the results of these control analyses are presented in supplementary materials only.

### 2.6. Lesion analyses

Univariate Voxel-based Lesion-Symptom Mapping (VLSM; Bates *et al*., 2003; Rorden *et al*., 2007) was used to identify anatomical regions associated with (1) disownership scores obtained during the baseline (pre-touch) condition of the experimental task (n = 24), and (2) ownership change scores obtained from the other-pleasant touch condition (n = 18). These VLSM analyses, when strictly corrected for multiple comparisons and minimum statistical power, did not show any significant results in our sample, possibly due to the small sample sizes, large lesions, and the fact that our experiment was optimised for subjective ownership changes in acute patients at the behavioural level (hence our main dependent variable had a relatively small variance range). In exploratory analyses we, therefore, repeated these analyses with less restrictive criteria (with no minimum lesion overlap and using 1% False Discovery Rate correction for multiple comparisons). We report in the main results below (section 3.3) only the significant findings from the exploratory analyses. Full details of the methods and results for these lesion analyses are reported in the supplementary materials.

### 2.7. Data availability

The data that support the findings of this study are available from the corresponding author, upon reasonable request.

## 3. Results

### 3.1. Clinical and neuropsychological results

The clinical and neuropsychological characteristics of patients are summarised in Table 1. Patients did not differ significantly in their clinical or neuropsychological profile. Importantly, even though the rNSA assessment of somatosensory function indicated that the perception of light touch was impaired in a number of patients, their ability to perceive both the pleasantness and intensity of repeated dynamic touch to some degree was confirmed in our control task that included sham trials to control for tactile perception (see Supplementary Materials).

### 3.2. Ownership change following touch

There was considerable variability in the extent to which each type of touch led to a change in ownership (see Figure 2). Despite this variability, a series of one-sample Wilcoxon signed-rank tests (with Bonferroni-corrected *α*=0.125 and pairwise deletion of cases with missing data) indicated that body ownership increased significantly following other affective touch (V(18) = 71, p = .012), but not following other types of touch (self-affective: V(20) = 53.50, p = 0.268; self-neutral: V(20) = 67, p = 0.371; other-neutral: V(20) = 65, p = 0.176; Figure 2). This analysis was also run using listwise deletion for missing data and Bayesian statistics (full details given in Supplementary Materials, section 3.1), confirming a substantive increase in body ownership following other-affective touch (BF_10_ = 6.95), but not other types of touch (all other BF_10_ > 0.3 < 3.0).

**Figure 2.**
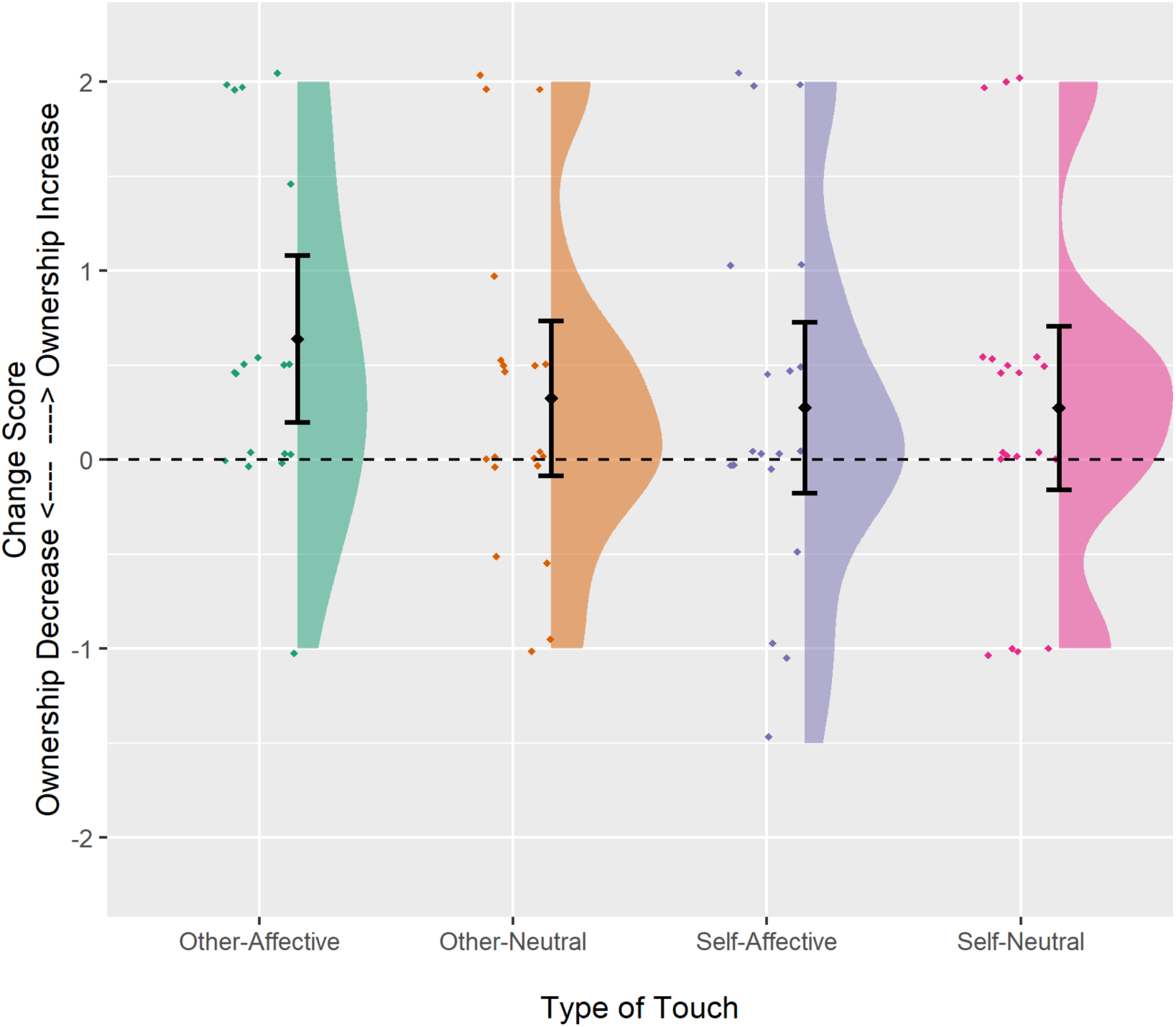
Raincloud plot illustrating change in body (dis)ownership after touch. Zero (dashed black line) indicates no change in (dis)ownership. Positive values indicate increased ownership/reduced DSO and negative values poorer ownership/increased DSO post-touch. The ‘cloud’ illustrates overall data distribution. Individual raw data are represented by the ‘rain’, with randomised jitter to improve visibility. Mean (black diamond) and 95% confidence interval (error bars) shown.

In addition to the formal assessment of ownership change reported above, during post-experiment debriefing we discussed any changes in body ownership with patients clinically, in order to understand the nature of their experience. One patient, who used to call her left arm her “alienated arm” in the first days following her stroke told us that, after the affective touch experiment, she would use her right, intact arm to stroke her left arm and speak to it; she said “come, correspond to me”. Another patient, somewhat similarly, told us a few days after the experiment: “I woke up and I called this [left] arm ‘a beast’. It was not my arm, I did not want it, it was some foreign fellow. But then you touched it and I caress it as you said and I decided to love it again. I said ‘Come, I accept you. I welcome you back’…We have been through a lot together”.

### 3.3. Lesion analyses

Our first exploratory lesion analysis, examining brain damage associated with baseline levels of DSO during the experimental session did not reveal any significant results (reported in Supplementary Materials only, for brevity). The second analysis, of lesions associated with a failure to increase limb ownership following other-affective touch, indicated that damage to the right insula, and more substantially the right corpus callosum, was associated with a failure for body ownership to increase following other-affective touch (FDR-corrected p < .01 for z > 2.42; see Figure 3. Full details of both analyses are provided in Supplementary Materials).

**Figure 3.**
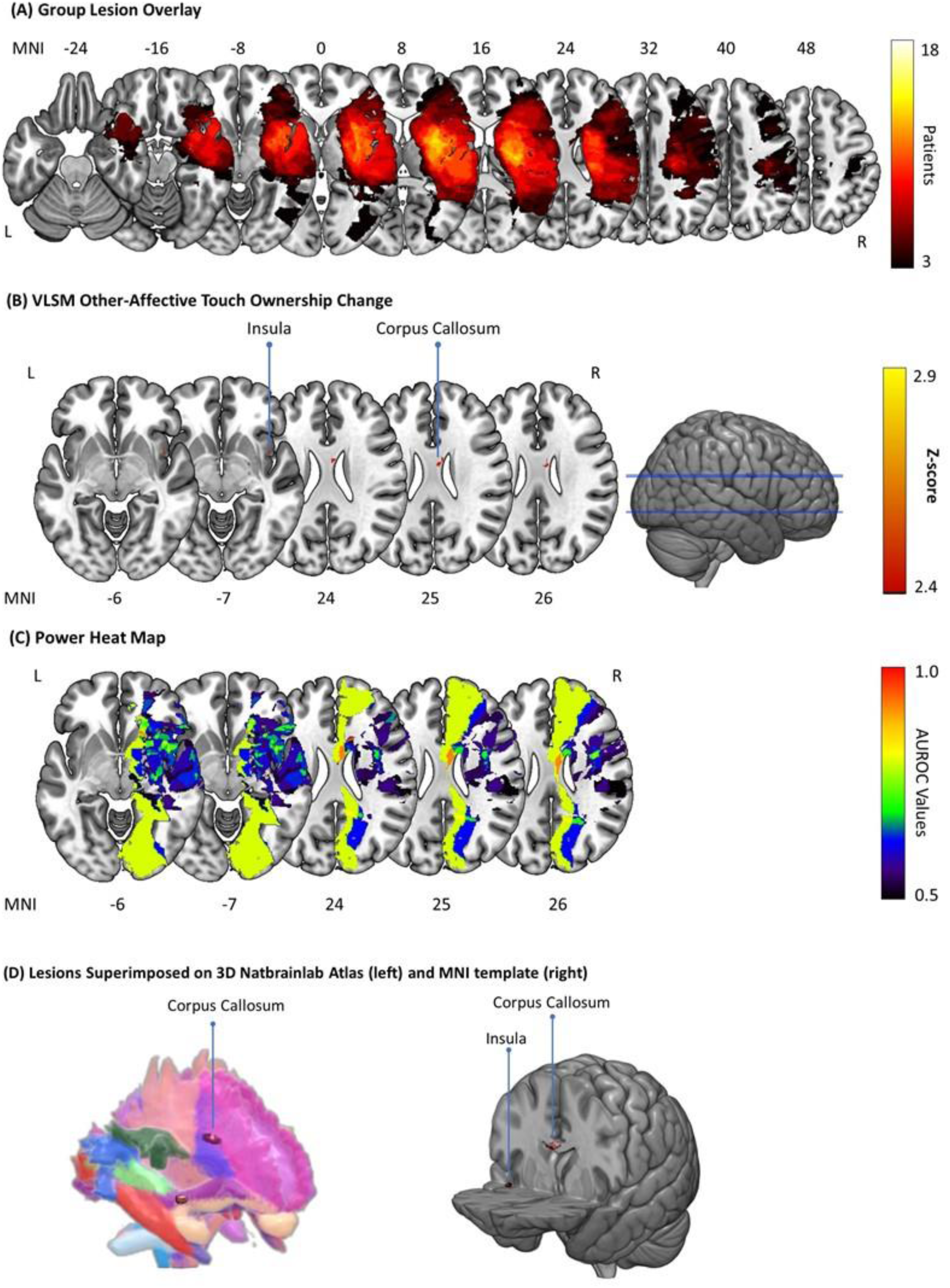
Lesion analysis. (A) Group-level lesion overlay maps. The number of overlapping lesions is illustrated from black (n=3) to white (n=18). (B) Damaged MNI voxels predicting failure to increase body ownership following other-affective touch (p < .01 for z > 2.42 FDR-corrected). (C) Heat map (AUROC) of the voxels with enough power to detect a significant result ranging between 0.5 (minimum power; shown in blue-black) to 1 (maximum discrimination power; shown in red). (D) 3D reproduction of lesions superimposed onto the Natbrainlab atlas (left) and MNI template (right).

## 4. Discussion

This study shows for the first time that receiving CT-optimal, affective touch can increase the sense of arm ownership in right-hemisphere stroke patients with DSO. These results are consistent with a growing body of experimental research in healthy subjects, which shows that affective, CT-optimal touch enhances feelings of body-part ownership during multisensory body-ownership illusions (Crucianelli *et al*., 2013, 2017; Lloyd *et al*., 2013; van Stralen *et al*., 2014; Panagiotopoulou *et al*., 2017; Ponzo *et al*., 2018). Below we consider the mechanisms by which CT-optimal touch administered by another person increases body ownership.

First, affective touch may reduce the unusual, sensory symptoms often reported by right hemisphere stroke patients, in particular “feelings of deafference”. We have previously argued that DSO may be the result of an inability to integrate current sensory signals with prior expectations about the sensory state of the body (Martinaud *et al*., 2017). Hence, disownership may occur when patients are unable to use current, aberrant signals from the body (e.g. feelings of ‘heaviness’, ‘numbness’, ‘coldness’ and other similar sensations) to update predictions and expectations about how the affected body parts should feel (Besharati and Fotopoulou, in press). This explanation is consistent with predictive coding accounts of body ownership, which propose that body ownership involves top-down, cross-modal predictions about the most likely cause of one’s bodily experience (e.g. the self) and requires the integration of exteroceptive, proprioceptive, and interoceptive sensory information (see Seth, 2013; Apps and Tsakiris, 2014; Fotopoulou, 2015; for further details and discussion). Indeed, research in healthy subjects using the rubber hand illusion shows that a temporal mismatch between seen and felt tactile stimulation (i.e. asynchronous, unpredictable touch) can produce several, unusual “deafference” sensations similar to those commonly reported by patients with DSO, including pins-and-needs, numbness and the hand being experienced as less vivid (Longo *et al*., 2008; Lesur *et al*., 2019). CT-optimal, affective touch, which provides an additional, cross-modal means of integrating different signals across modalities (namely, “affective congruency”; see Filippetti *et al*., 2019 for further discussion) and has been found to reduce “deafference” in healthy subjects (see Panagiotopoulou *et al*., 2017), may similarly lessen feelings of deafference in patients with DSO.

Affective touch may also increase arm ownership by enhancing interoceptive signals known to play an important role in the “feeling of mineness” (see Craig, 2003; Crucianelli *et al*., 2017). Interoception informs the mind about how the body is doing in relation to certain homeostatic needs (e.g. hunger, thirst, pain and pleasure), and is considered by some modern neuroscientific theories to be the basis of subjective feeling states and the sentient self (Craig, 2009; Seth, 2013; Fotopoulou and Tsakiris, 2017). Thus, in the present study CT-optimal touch may change beliefs about body ownership by strengthening these fundamental feelings of mineness. This effect is specific to the touched limb, and not a general valence effect, as demonstrated by the fact that applying affective touch to the unaffected right arm did not increase ownership of the affected left arm.

Finally, touch from another person may draw attention to the left hand, thereby enhancing the salience (i.e. the epistemic importance of the hand in relation to other percepts) of signals arising from the affected limb. An increase in attention might result in signals from the affected limb being given greater weight during multisensory integration – which can produce an update in knowledge about the state of the body and change in beliefs about limb ownership (see e.g. Bolognini *et al*., 2014; D’Imperio *et al*., 2017). In the context of this explanation, touch from another person might be deemed more salient owing to its lack of concurrent efferent information and consequently greater unpredictability (Blakemore *et al*., 1998*b*, 1999). However, our results suggest that enhanced salience or attention is unlikely to be the main mechanism of change in our patients, since visuo-spatial and personal neglect for the affected side and arm were not consistently or concurrently increased by affective touch; CT-optimal touch led to an increase specifically in arm ownership.

Our supplementary, exploratory analyses also indicated that not all patients were able to perceive touch reliably, or the difference between affective and neutral touch intensity and pleasantness. There was also no relationship between the perceived intensity or pleasantness of touch and baseline disownership or the ownership change. Earlier reports found that self-touch enhances the perception of touch in a small group of right-hemisphere patients (White *et al*., 2010). Our own exploratory examination of the perceived intensity and pleasantness of touch revealed that the *intensity* of self-touch was greater than that of other-touch (consistent with White and colleagues), irrespective of velocity, while the perceived *pleasantness* of touch did not differ for self-versus other-touch or different velocities. Thus, our findings show that even when CT-optimal touch received from an “other” is perceived as less intense and as equally pleasant as self-touch, it can be particularly potent to increase limb ownership. This finding supports the observed dissociation between neural systems responsible for discriminative versus affective aspects of touch (see McGlone *et al*., 2014 for a review), and suggests that this dissociation includes the selective enhancement of discriminative self-touch, while not affecting perception of the CT-touch system. However, these findings need to be interpreted with caution as we had limited measurements of patients’ tactile perception, either by standardised assessments or, during the task (only four trials per touch type). Future studies should explore CT touch effects in association with systematic, somatosensory assessments.

Our study also has important implications for the rehabilitation of patients with typically poor therapy adherence and engagement. A simple-to-administer, non-verbal, non-invasive affective touch procedure requiring no specialist training or equipment can provide an effective means of improving post-stroke DSO. Despite our averaged findings and the consistent spontaneous remarks from some of our patients, we note that not all patients with DSO showed increased ownership following other-affective touch, and we did not include follow-ups in the present, experimental study. More generally, further research is needed to identify minimum, critical factors that determine who can benefit from affective touch, and to establish the optimum dose (frequency, timing and duration of treatment), extent and stability of effects. Additionally, further work is needed to explore factors that might moderate the efficacy of affective touch, such as the ability to perceive touch and affective touch, baseline degree of disownership, and concurrent neuropsychological deficits.

One potentially fruitful avenue to explore is the identification of neuroanatomical predictors (Lunven *et al*., 2015; Forkel and Catani, 2018). Our initial lesion analyses did not yield any significant results, likely due to the small sample, large lesions and limited range of scores possible on our behavioural measure. However, our exploratory VLSM using less restrictive criteria revealed that lesions in the insula and corpus callosum, and, resulted in significantly less increase in body ownership following other-affective touch. These findings are consistent with the proposed role of the insula in body awareness, as well as the importance of white matter tracts in self-awareness (Pacella *et al*., 2019) and body ownership (Moro *et al*., 2016; Martinaud *et al*., 2017) and warrant further study.

In conclusion, we report novel findings from right hemisphere stroke patients with disturbances in body ownership. Using simple tactile stimulation parameters that are known to activate the CT system optimally, we were able to increase the sense of arm ownership in patients with DSO. Based on a number of additional manipulation, we suggest that these interoceptive signals bring about change in ownership by reducing feelings of deafference and allowing new sensations from the affected body parts to be integrated with one’s multi-modal, self-representation, rather than less likely mechanisms of attentional enhancement. Finally, our study provides the first experimental evidence for the use of neurophysiologically-specified type of affective touch in the treatment of DSO that can be further tested in translational studies.

## Acknowledgements

We thank the patients and their relatives for their kindness and willingness to take part in the study, and to the staff at participating hospital for their assistance with this study. We are also grateful to Dr Sonia Ponzo for her help with patient recruitment and testing, and to Dr Thanos Koukoutsakis for creating the raincloud plots used in the paper.

## Author contributions

PMJ = Paul M Jenkinson, CP = Cristina Papadaki, Sahba Besharati, = SB, Valentina Moro = VM, Valeria Gobbetto = VG, Laura Crucianelli = LC, Louise P Kirsch = LPK, Renato Avesani = RA, Nicholas Ward = NW, Aikaterini Fotopoulou = AF.

Study conception: PMJ, CP, SB, LC, AF.

Methodology (design): PMJ, CP, SB, LC, AF.

Investigation (data collection): SB, LC, CP, LPK, VG.

Investigation (lesion drawing): VM.

Analysis: PMJ.

Data curation: SB, CP, LPK.

Resources: VM, NW, AF.

Writing (initial draft): PMJ, AF.

Writing (critical review, commentary or revision): PMJ, SB, CP, LC, VM, NW, LPK, AF.

Supervision: PMJ, VM, AF.

Project administration: PMJ, VM, AF, LPK.

Funding acquisition: VM, AF.

## References

Ackerley R, Hassan E, Curran A, Wessberg J, Olausson H, McGlone F. An fMRI study oncortical responses during active self-touch and passive touch from others. Front Behav Neurosci 2012; 6: 1–9.

Apps MAJ, Tsakiris M. The free-energy self: A predictive coding account of self-recognition. Neurosci Biobehav Rev 2014; 41: 85–97.

Baier B, Karnath H-O. Tight link between our sense of limb ownership and self-awareness of actions. Stroke 2008; 39: 486–8.

Bates E, Wilson SM, Saygin AP, Dick F, Sereno MI, Knight RT, et al. Voxel-based lesion-symptom mapping. Nat Neurosci 2003; 6: 448–450.

Besharati S, Crucianelli L, Fotopoulou A. Restoring awareness: a review of rehabilitation in anosognosia for hemiplegia. Rev Chil Neuropsicol 2014; 9: 31–37.

Besharati S, Fotopoulou A. The social reality of the self: Right perisylvian damage revisited. In: Salas C, Turnbull O, Solms M, editor(s). Clinical studies in neuropsychoanalysis revisited. Oxford, UK: Routledge

Besharati S, Kopelman M, Avesani R, Moro V, Fotopoulou AK. Another perspective on anosognosia: Self-observation in video replay improves motor awareness. Neuropsychol Rehabil 2014: 1–34.

Bisiach E, Vallar G, Perani D, Papagno C, Berti A. Unawareness of disease following lesions of the right hemisphere: Anosognosia for hemiplegia and anosognosia for hemianopia. Neuropsychologia 1986; 24: 471–482.

Blakemore S-J, Wolpert DM, Frith CD. Central cancellation of self-produced tickle sensation. Nat Neurosci 1998; 1: 635–640.

Blakemore SJ, Frith CD, Wolpert DM. Spatio-temporal prediction modulates the perception of self-produced stimuli. J Cogn Neurosci 1999; 11: 551–9.

Blakemore SJ, Goodbody SJ, Wolpert DM. Predicting the Consequences of Our Own Actions: The Role of Sensorimotor Context Estimation. J Neurosci 1998; 18: 7511–7518.

Boehme R, Hauser S, Gerling GJ, Heilig M, Olausson H. Distinction of self-produced touch and social touch at cortical and spinal cord levels. Proc Natl Acad Sci U S A 2019; 116: 2290–2299.

Bolognini N, Ronchi R, Casati C, Fortis P, Vallar G. Multisensory remission of somatoparaphrenic delusion: My hand is back! Neurol Clin Pract 2014; 4: 216–225.

Botvinick M, Cohen J. Rubber hands ‘feel’ touch that eyes see. Nature 1998; 391: 756–756.

Case LK, Laubacher CM, Olausson H, Wang B, Spagnolo PA, Bushnell MC. Encoding of Touch Intensity But Not Pleasantness in Human Primary Somatosensory Cortex. J Neurosci 2016; 36: 5850–5860.

Ceunen E, Vlaeyen JWS, Van Diest I. On the origin of interoception. Front Psychol 2016; 7: 1–17.

Craig A. Interoception: the sense of the physiological condition of the body. Curr Opin Neurobiol 2003; 13: 500–505.

Craig AD. How do you feel - now? The anterior insula and human awareness. Nat Rev Neurosci 2009; 10: 59–70.

Crucianelli L, Krahé C, Jenkinson PM, Fotopoulou A. Interoceptive ingredients of body ownership: Affective touch and cardiac awareness in the rubber hand illusion. Cortex 2017

Crucianelli L, Metcalf NKNK, Fotopoulou AK, Jenkinson PMPM. Bodily pleasure matters: velocity of touch modulates body ownership during the rubber hand illusion. Front Psychol 2013; 4: 703.

Crucianelli L, Paloyelis Y, Ricciardi L, Jenkinson PM, Fotopoulou A. Embodied Precision: Intranasal Oxytocin Modulates Multisensory Integration. J Cogn Neurosci 2019; 31: 592–606.

D’Imperio D, Tomelleri G, Moretto G, Moro V. Modulation of somatoparaphrenia following left-hemisphere damage. Neurocase 2017; 23: 162–170.

Dienes Z. Using Bayes to get the most out of non-significant results. Front Psychol 2014; 5: 1–17.

Dienes Z, Mclatchie N. Four reasons to prefer Bayesian analyses over significance testing. Psychon Bull Rev 2018; 25: 207–218.

Dubois B, Slachevsky A, Litvan I, Pillon B. The FAB: A frontal assessment battery at bedside. Neurology 2000; 55: 1621 LP –1626.

Feinberg TE, Roane DM, Ali J. Illusory limb movements in anosognosia for hemiplegia. J Neurol Neurosurg Psychiatry 2000; 68: 511–3.

Filippetti ML, Kirsch LP, Crucianelli L, Fotopoulou A. Affective certainty and congruency of touch modulate the experience of the rubber hand illusion. Sci Rep 2019; 9: 1–13.

Forkel SJ, Catani M. Lesion mapping in acute stroke aphasia and its implications for recovery. Neuropsychologia 2018; 115: 88–100.

Fotopoulou A. The virtual bodily self: Mentalisation of the body as revealed in anosognosia for hemiplegia. Conscious Cogn 2015; 33: 500–510.

Fotopoulou A, Tsakiris M. Mentalizing homeostasis : The social origins of interoceptive inference. 2017; 4145

Gentsch A, Panagiotopoulou E, Fotopoulou A. Active Interpersonal Touch Gives Rise to the Social Softness Illusion. Curr Biol 2015; 25: 2392–2397.

Guarantors of Brain. Aids to the examination of the peripheral nervous system. London: W. B. Saunders; 1986

Halligan PW, Cockburn J, Wilson BA. The behavioural assessment of visual neglect. Neuropsychol Rehabil 1991; 1: 5–32.

Howard GS, Maxwell SE, Fleming KJ. The proof of the pudding: An illustration of the relative strengths of null hypothesis, meta-analysis, and bayesian analysis. Psychol Methods 2000; 5: 315–332.

JASP Team. JASP (Version 0.11.1)[Computer software]. 2019

Jeffreys H. The Theory of Probability. 3rd ed. Oxford, UK: Oxford University Press; 1961

Jenkinson PM, Moro V, Fotopoulou A. Definition: Asomatognosia. Cortex 2018; 101: 300–301.

Jenkinson PM, Preston C, Ellis SJ. Unawareness after stroke: A review and practical guide to understanding, assessing, and managing anosognosia for hemiplegia. J Clin Exp Neuropsychol 2011; 33: 1079–1093.

Kilteni K, Ehrsson HH. Body ownership determines the attenuation of self-generated tactile sensations. Proc Natl Acad Sci 2017; 114: 8426–8431.

Lesur MR, Weijs ML, Simon C, Kannape OA, Lenggenhager B. Achronopresence: how temporal visuotactile and visuomotor mismatches modulate embodiment. bioRxiv 2019: 596858.

Lincoln NB, Jackson JM, Adams SA. Reliability and revision of the Nottingham Sensory Assessment for stroke patients. Physiotherapy 1998; 84: 358–365.

Lloyd DM, Gillis V, Lewis E, Farrell MJ, Morrison I. Pleasant touch moderates the subjective but not objective aspects of body perception. Front Behav Neurosci 2013; 7: 207.

Löken LS, Wessberg J, Morrison I, McGlone F, Olausson H. Coding of pleasant touch by unmyelinated afferents in humans. Nat Neurosci 2009; 12: 547–8.

Longo MR, Schüür F, Kammers MPM, Tsakiris M, Haggard P. What is embodiment? A psychometric approach. Cognition 2008; 107: 978–998.

Lunven M, Thiebaut De Schotten M, Bourlon C, Duret C, Migliaccio R, Rode G, et al. White matter lesional predictors of chronic visual neglect: a longitudinal study. Brain 2015: 1–15.

Martinaud O, Besharati S, Jenkinson PM, Fotopoulou A. Ownership illusions in patients with body delusions: Different neural profiles of visual capture and disownership. Cortex 2017; 87

Mcglone F, Olausson H, Boyle JA, Jones-Gotman M, Dancer C, Guest S, et al. Touching and feeling: Differences in pleasant touch processing between glabrous and hairy skin in humans. Eur J Neurosci 2012; 35: 1782–1788.

McGlone F, Wessberg J, Olausson H. Discriminative and Affective Touch: Sensing and Feeling. Neuron 2014; 82: 737–755.

McIntosh RD, Brodie EE, Beschin N, Robertson IH. Improving the Clinical Diagnosis of Personal Neglect: A Reformulated Comb and Razor Test. Cortex 2000; 36: 289–292.

Moro V, Pernigo S, Tsakiris M, Avesani R, Edelstyn NMJ, Jenkinson PM, et al. Motor versus body awareness: Voxel-based lesion analysis in anosognosia for hemiplegia and somatoparaphrenia following right hemisphere stroke. Cortex 2016; 83: 62–77.

Moro V, Zampini M, Aglioti SM. Changes in Spatial Position of Hands Modify Tactile Extinction but not Disownership of Contralesional Hand in Two Right Brain-Damaged Patients. Neurocase 2004; 10: 437–443.

Morrison I. ALE meta-analysis reveals dissociable networks for affective and discriminative aspects of touch. Hum Brain Mapp 2016

Morrison I, Löken LS, Minde J, Wessberg J, Perini I, Nennesmo I, et al. Reduced C-afferent fibre density affects perceived pleasantness and empathy for touch. Brain 2011; 134: 1116–1126.

Nasreddine ZS, Phillips NA, Bédirian V, Charbonneau S, Whitehead V, Collin I, et al. The Montreal Cognitive Assessment, MoCA: A Brief Screening Tool For Mild Cognitive Impairment. J Am Geriatr Soc 2005; 53: 695–699.

Nordin M. Low-threshold mechanoreceptive and nociceptive units with unmyelinated (C) fibres in the human supraorbital nerve. J Physiol 1990; 426: 229–240.

Olausson H, Lamarre Y, Backlund H, Morin C, Wallin BG, Starck G, et al. Unmyelinated tactile afferents signal touch and project to insular cortex. Nat Neurosci 2002; 5: 900–904.

Pacella V, Foulon C, Jenkinson PM, Scandola M, Bertagnoli S, Avesani R, et al. Anosognosia for hemiplegia as a tripartite disconnection syndrome [Internet]. Elife 2019; 8 Available from: https://elifesciences.org/articles/46075

Panagiotopoulou E, Filippetti ML, Tsakiris M, Fotopoulou A. Affective Touch Enhances Self-Face Recognition during Multisensory Integration. Sci Rep 2017; 7: 1–10.

Ponzo S, Kirsch LP, Fotopoulou A, Jenkinson PM. Balancing body ownership: Visual capture of proprioception and affectivity during vestibular stimulation. Neuropsychologia 2018; 117: 311–321.

Quintana DS, Williams DR. Bayesian alternatives for common null-hypothesis significance tests in psychiatry: A non-technical guide using JASP. BMC Psychiatry 2018; 18: 1–8.

R Core Team. R: A Language and Environment for Statistical Computing [Internet]. 2013 Available from: http://www.r-project.org/

Rorden C, Brett M. Stereotaxic display of brain lesions. Behav Neurol 2000; 12: 191–200.

Rorden C, Karnath H-O, Bonilha L. Improving Lesion-Symptom Mapping. J Cogn Neurosci 2007; 19: 1081–1088.

Seth AK. Interoceptive inference, emotion, and the embodied self. Trends Cogn Sci 2013; 17: 565–73.

van Stralen HE, van Zandvoort MJE, Hoppenbrouwers SS, Vissers LMG, Kappelle LJ, Dijkerman HC. Affective touch modulates the rubber hand illusion. Cognition 2014; 131: 147–58.

van Stralen HE, van Zandvoort MJEE, Dijkerman HC. The role of self-touch in somatosensory and body representation disorders after stroke. Philos Trans R Soc B Biol Sci 2011; 366: 3142–3152.

Valentini M, Kischka U, Halligan PW. Residual haptic sensation following stroke using ipsilateral stimulation. J Neurol Neurosurg Psychiatry 2008; 79: 266–270.

Vallar G, Ronchi R. Somatoparaphrenia: a body delusion. A review of the neuropsychological literature. Exp Brain Res 2009; 192: 533–551.

Vallbo Å, Olausson H, Wessberg J, Norrsell U. A system of unmyelinated afferents for innocuous mechanoreception in the human skin. Brain Res 1993; 628: 301–304.

Vallbo ÅB, Hagbarth KE. Activity from skin mechanoreceptors recorded percutaneously in awake human subjects. Exp Neurol 1968; 21: 270–289.

Vallbo ÅB, Olausson H, Wessberg J. Unmyelinated afferents constitute a second system coding tactile stimuli of the human hairy skin. J Neurophysiol 1999; 81: 2753–2763.

Wechsler D. WAIS-III: administration and scoring manual: Wechsler adult intelligence scale. 1997

Wechsler D. Wechsler test of adult reading: WTAR. 2001

White RC, Davies AMA, Kischka U, Davies M. Touch and feel? Using the rubber hand paradigm to investigate self-touch enhancement in right-hemisphere stroke patients. Neuropsychologia 2010; 48: 26–37.

Wickham H. ggplot2: Elegant Graphics for Data Analysis [Internet]. 2016 Available from: https://ggplot2.tidyverse.org

Zigmond AS, Snaith RP. The Hospital Anxiety and Depression Scale. Acta Psychiatr Scand 1983

