## Supplementary for "Welcoming back my arm: Affective touch increases body ownership following right hemisphere stroke"

### Supplementary Materials

### 1. Feasibility study

In order to inform the procedures and predictions of the main experiment, and particularly the feasibility of applying and measuring the effect of repeated touch stimuli to this population, including the acceptability, fatigue, concentration and comprehension of patients, we conducted an extensive feasibility / pilot study with 16 right hemisphere patients recruited using the same inclusion/exclusion criteria as the main study. This study provided feedback on overall patient attitude towards the tactile protocol, and optimisation of the measurement scale, but was not able to inform the hypothesis of the main study regarding the potentially specific role of affective vs. neutral touch on body ownership. Instead, we measured the overall, emotional attitude and feelings of limb ownership only before and after the entire touch protocol which included both affective (CT-optimal: 3cm/s) and neutral (CT-suboptimal: 18cm/s) tactile stimulation.

We initially assessed the ability of patients to understand different variations of the rating scale (e.g. horizontal vs. vertical presentation, with/without numerical descriptors, and with greater/ fewer anchor words). We then assessed their ability to follow instructions, detect, discriminate and tolerate repeated trials of slow and fast stroking, applied to the left and right arm, as well as sham trials (during which slow velocity stroking was simulated just above the arm without applying actual touch), and a non-hairy-skin control condition. Following each trial patients were asked whether they had perceived the touch ("did you feel touch? Yes or no?"), where ("on which arm? Left or right?"), and its pleasantness ("how pleasant/unpleasant?" 0 = not at all to 5 = extremely pleasant). Arm ownership was assessed before and after the entire touch procedure by asking four questions: “Is this your hand? Does it ever feel like it does not belong to you? Do you ever feel that your arm is missing?” Do you ever feel that your arm has disappeared?”, which were scored using the same scale as the main study (0.5 = no disownership; 0.5 = partial disownership; 1 = disownership) and summed to give a score ranging from 0 (minimal disownership) to 4 (maximum disownership).

We found that patients were able to detect and discriminate between fast CT-suboptimal and slow CT-optimal touch as demonstrated in their ratings (reported fully in a separate study by Kirsch *et al.*, 2019) and qualitative comments: a number of patients remarked that the slow touch elicited a pleasant sensation, with one patient describing it as “light and soft” while the fast touch was “rough”. Another patient, upon being touched using the slow velocity, spontaneously laughed and said “this is ticklish! Quite pleasant though”. However, patients also commented on the protocol being long and repetitive, owing to multiple questions to which they gave the same response, and touch being applied many times. As a result, the ability to detect and discriminate touch was explored fully in a separate study (Kirsch *et al.*, 2019), while in further developing the main experiment reported here we focused on assessing ownership before and after each type of touch, by using a block design in which we reduced the number of touch trials (from 6 to 4) and sham (to one per block), combined and reduced the number of ownership questions, and performed any control conditions in separate blocks to avoid boredom, frustration and fatigue.

At the group level, ownership (calculated using the formula: pre-ownership score – post-ownership score) was significant increased by touch as indicated by a one-sample Wilcoxon signed-rank test ($V(11) = 21$, $p = .036$; Fig. S1), and confirmed by a Bayesian one-sample t-tests ($BF_{10} = 3.17$; see section 2.5 of main document for details of Bayesian analysis and interpretation). We also noted several interesting remarks from patients regarding their ownership of the paralysed arm. For example, a patient with somatoparaphrenia showed a striking increase in arm ownership following the touch procedure. Before the touch the patient consistently denied ownership of the left hand, claiming that it belonged to a friend, felt like it did not belong to her and was missing. Post-touch the patient recognised the arm as her own. Asked if it ever felt like it did not belong, she replied “well now, no”, and when asked if she ever felt like the arm was missing she said “not really”. Another patient with pre-touch asomatognosia (feelings of non-belonging towards the arm), showed an increase in arm ownership, and when asked if the arm ever felt like it did not belong to her post-stroking, stated “not too much now, the brushing helped me”.

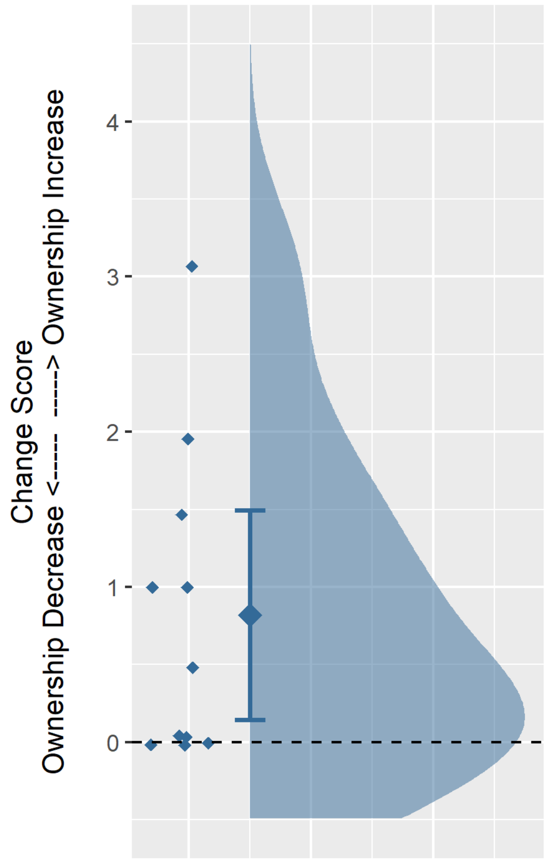

***Figure S1.*** *Mean ownership change following touch in 11 patients showing baseline disownership. Dashed line (zero) indicates no change in ownership. A positive value indicates an increase in ownership post-touch. Overall distribution of scores represented by*

*the 'cloud'. Raw data indicated by the 'rain' with randomized jitter for improved visualisation. Error bars represent 95% confidence interval around the mean (central diamond).*

### 2. Main Experiment - Control Tasks Methods

In order to further explore the specificity and mechanism of the touch effect, within practical testing constraints, we conducted a number of control assessments and analyses in a small subset of patients.

Firstly, we assessed left-sided visuospatial and personal neglect before and after each affective touch condition using the 'copy' star and flower subtest of the BIT (Halligan *et al.*, 1991), and the one-item test (Bisiach *et al.*, 1986). For the copy task, we scored each drawing as: 0 = majority of figure omitted, 0.5 = partial completion with some omission, 1 = no omission / complete drawing. The one-item test was administered by asking the patient to reach and touch their contralesional (left) hand with their ipsilesional (right) hand while their eyes were closed, and was scored by the experimenter: 0 = the patient promptly reaches for the target, 1 = the target is reached with hesitation and search, 2 = the search is interrupted before the target is reached, 3 = no movement towards the target is performed. Following each task, a measure of neglect awareness was obtained by asking the patient to rate how well they had performed the task using a scale from zero (not well at all) to ten (very well).

Secondly, we applied self- (n=16) and other-affective (n=15) touch to the *right* arm of patients using the same procedure as for the left arm, and assessed any changes in ownership regarding the *left* (disowned) limb. This control condition was performed after the four main touch conditions in the majority of patients (n self = 15; n other = 12); however, to ensure that any observed effects of this condition were not simply an order effect, we also randomly varied the position of the two right arm blocks such that they occurred amongst the four main conditions in a subset of patients (n = 5).

Finally, although time and fatigue considerations prevented us from using established protocols for affective touch perception and discrimination (Gentsch *et al.*, 2015; von Mohr *et al.*, 2019) in order to evaluate the effect of touch perception on ownership change, we did collect some subjective ratings. Patients were asked to rate the intensity (i.e. “how well did you feel the touch?”) of each touch trial using a vertical scale (to counteract the influence of any visuospatial neglect) ranging from 0 (“not at all”) to 10 (“very well”). Subsequently, patients rated the pleasantness of each touch trial (i.e. “how pleasant was the touch?”) using a 6-point, vertical scale ranging from “not at all pleasant” (scored 0) to “extremely pleasant” (scored 5). If the patients did not report any touch on a given trial (i.e. intensity = 0/not at all) they were not asked to provide a pleasantness rating, and the average touch pleasantness was calculated using the completed trials. We chose to use two different scales for the intensity and pleasantness ratings to clearly differentiate between the two ratings (intensity vs. touch), and reduce perseveration and response repetition.

### 3. Main Experiment - Control Tasks and Order Effects Analyses

**Order effects**: as detailed above, we were unable to fully counterbalance the order of experimental touch blocks; therefore, in post-hoc, supplementary analyses we examined whether there was a significant bias in the order frequency of the four main conditions. In order to do this we calculated the frequency with which each condition appeared in each testing position (i.e. 1st, 2nd, 3rd or 4th; see Supplementary Table S1), and subsequently used the chi-squared goodness-of-fit test to assess if there was a significant deviation from an expected, even distribution of block orders (i.e. a significant result would indicate that some touch blocks occurred more frequently in a particular order position). In addition, because the other affective touch condition was the only one to produce a reliable increase in body ownership (see main results), we conducted a follow-up analyses to explore if this increase was affected by its position amongst the four main conditions (using a Kruskall-Wallis test), or being located in the first two versus latter two touch conditions (Mann-Whitney *U* test).

**Neglect:** to determine the specificity of the affective touch effect, and whether this might be related to a more general change in visuospatial or body-related attention, we examined whether affective touch changed the degree of visuospatial and personal neglect in two DSO patients. We report the pattern of change in these cases descriptively, since statistical analysis is not possible with so few cases.

**Right hand:** to determine if applying touch to the right hand affected disownership of the left hand, we ran the same one-sample tests described for the main analysis, but using the ratings obtained after stroking the right hand to examine if touch caused a change in ownership greater than zero. Additionally, to confirm the specificity of our finding regarding other-affective touch on the left hand, we directly compared within-subject whether or not other-affective touch applied to the left versus right hand produced an increase in left arm ownership, using the Exact McNemar's test for binomial data. To perform this analysis we categorised patients in terms of whether or not their left arm ownership increased as a result of other-affective touch to the left versus right arm. We did not expect to see any significant change in disownership of the left hand as a result of the right hand touch conditions, but expected other-affective touch on the left hand to cause a significant increase in ownership relative to the same touch applied to the right hand.

**Touch pleasantness and intensity:** we examined if the degree of baseline disownership (DSO) and disownership change was related to the perception of touch on the left arm. The average intensity and pleasantness score (measured after each of the four trials of each touch type/condition) was calculated for each DSO patient. The effect of touch velocity (slow vs. fast) and agent (self vs. other) on these intensity and pleasantness ratings was then examined, followed by the relationship between these two aspects of touch perception and (1) body ownership at baseline as well as (2) ownership change scores, using bivariate, non-parametric (Spearman's) and Bayesian correlations.

### 4. Main Experiment - Supplementary Results

#### 4.1. Ownership Change – Listwise deletion and Bayesian analysis

The main experimental dependent variable was analysed using both pairwise deletion for missing data (reported in main manuscript) and listwise deletion, to account for possible differences in the results arising from missing data and the use of these different methods. Results of the listwise deletion analysis confirmed that a significant increase in body ownership was present only after other-affective touch (V(18) = 71, p = .012), and not the other three types of touch: self-affective touch V (18) = 49.50, p = .151; self-neutral touch V (18) = 61, p = .084; other-neutral touch V(18) = 55, p = .051 (Bonferroni-corrected $\alpha$=.0125 for four comparisons).

Bayesian one-sample t-tests were likewise run using both pairwise and listwise deletion methods. Results of the pairwise Bayesian analysis showed support for the other-affective touch (n = 18) improving body ownership ($BF_{10}$ = 6.95), but not the other three types of touch (n = 20): self-affective $BF_{10}$ = .47; self-neutral $BF_{10}$ = .50; other-neutral $BF_{10}$ = .742. The same analysis done using listwise deletion (all n = 18) showed greater support for self-neutral and other-neutral touch improving body ownership, but these were not above recommended benchmarks ($BF_{10}$ >3 or <0.3) and provided only anecdotal evidence, which did not change the pattern of results: other-affective $BF_{10}$ = 6.952; self-affective $BF_{10}$ = .697; self-neutral $BF_{10}$ = 1.322; other-neutral $BF_{10}$ = 1.788.

#### 4.2. Order effects

To examine if order – and specifically the position of the other affective touch condition – affected the observed increase in body ownership, we undertook supplementary analyses of order effects (described above). These analyses showed that the other affective touch was administered more frequently as the second condition (44% of the time); however, a chi-square goodness of fit test indicated that the position of the other-affective condition was not significantly different from that expected by chance ( $\chi^2$ (3, N=18) = 3.28, p = .286). As further confirmation, we treated position as an independent variable and examined whether the effect of other affective touch on change in body ownership depended on its position (1st, 2nd, 3rd, 4th) in the order of conditions, using the Kruskall-Wallis test. Results indicated no significant effect of touch position on the effect of other affective touch (H (3) = 2.56, p = .465). However, because the Kruskall-Wallis test may have been influenced by the small sample and subsequently low statistical power, we also performed this analysis by classifying the position of the other affective touch condition as in the first half (i.e. position 1 or 2) or second half (i.e. position 3 or 4) and then comparing whether the effect of other affective touch on body ownership increase occurred more frequently in the first or second half of testing conditions, using the Mann-Whitney *U* test. Results again indicated no significant effect of position on the effect of other affective touch (U = 34, p = .705).

***Table S1.*** *Frequency with which the other-affective touch condition appeared in each position.*

| **Position** | **Frequency** | **% of total** |
|---|---|---|
| First | 3 | 16.7 |
| Second | 8 | 44.4 |

| | | |
|---|---|---|
| Third | 3 | 16.7 |
| Fourth | 4 | 22.2 |

**4.3. Specificity of effect: Neglect**

Two DSO patients completed the neglect and awareness of neglect assessments, the results of which are summarised in Supplementary Table S2. Overall, these individual cases did not show a consistent pattern of behaviour that indicated a robust effect of either self- or other-affective-touch on neglect or awareness of neglect. Prior to self-affective-touch, patient DSO16 exhibited mild extrapersonal, and personal neglect. There was a decrease in both extrapersonal and personal neglect after self-affective-touch, but no change in neglect awareness. There was no sign of (extrapersonal or personal) neglect prior to other-affective-touch, no change in neglect as a result of touch, and no change in neglect awareness (except for an increase in the patient's rating following other-affective-touch, despite no change in actual performance). By contrast, patient DSO17 presented with no extrapersonal or personal neglect prior to self-affective-touch and subsequently no change in neglect as a result of the touch. Mild extrapersonal neglect, but no personal neglect, was present prior to other-affective-touch. This extrapersonal neglect decreased after stroking, but awareness did not (in fact, ratings of ability to perform the task decreased slightly – despite better performance post-touch).

***Table S2.*** *Experimental results of additional control experiment with 2 right-hemisphere brain damaged patients (DSO16 & DSO17) investigating change in visuospatial neglect, personal neglect and awareness of neglect.*

| Modality & Test | Patient | Self affective touch | | Other affective touch | |
|---|---|---|---|---|---|
| | | Pre | Post | Pre | Post |
| **Extrapersonal neglect**<br>copy star / flower<br>(Range: 0-3) | | | | | |
| | DSO16 | 0.5 | 0 | 0 | 0 |
| | DSO17 | 0 | 0 | 0.5 | 0 |
| **Personal neglect**<br>One item Test<br>(Range: 0-3) | | | | | |
| | DSO16 | 2 | 0 | 0 | 0 |
| | DSO17 | 0 | 0 | 0 | 0 |
| **Awareness of extrapersonal neglect**<br>Patient rating<br>(Range: 0-10) | | | | | |
| | DSO16 | 9 | 9 | 4 | 9 |
| | DSO17 | 5 | - | 2 | 0 |
| **Awareness of personal neglect**<br>Patient rating<br>(Range: 0-10) | | | | | |
| | DSO16 | 10 | 10 | 10 | 10 |

| | | | | |
|---|---|---|---|---|
| DSO17 | 5 | - | 10 | 10 |

*Note: One item test: 0 = the patient promptly reaches for the target; 1 = the target is reached with hesitation and search; 2 = the search is interrupted before the target is reached; 3 = no movement towards the target is performed. Copy star / flower: 0 = majority of figure omitted, 0.5 = partial completion with some omission, 1 = no omission / complete drawing. Awareness rating scale for performance on copy / one item test: 0 = not well at all, 10 = very well. Dashed lines indicate missing data.*

#### 4.4. Specificity of effect: Right hand touch

Separate one-sample Wilcoxon signed-rank tests (with Bonferroni-corrected $\alpha$=.025 and pairwise deletion of cases with missing data) indicated that touch to the right hand did not produce a significant change in disownership (self-affective touch: $V(16) = 72$, $p = .227$; other-affective touch: $V(15) = 49.50$, $p = .150$), see Figure S1 below.

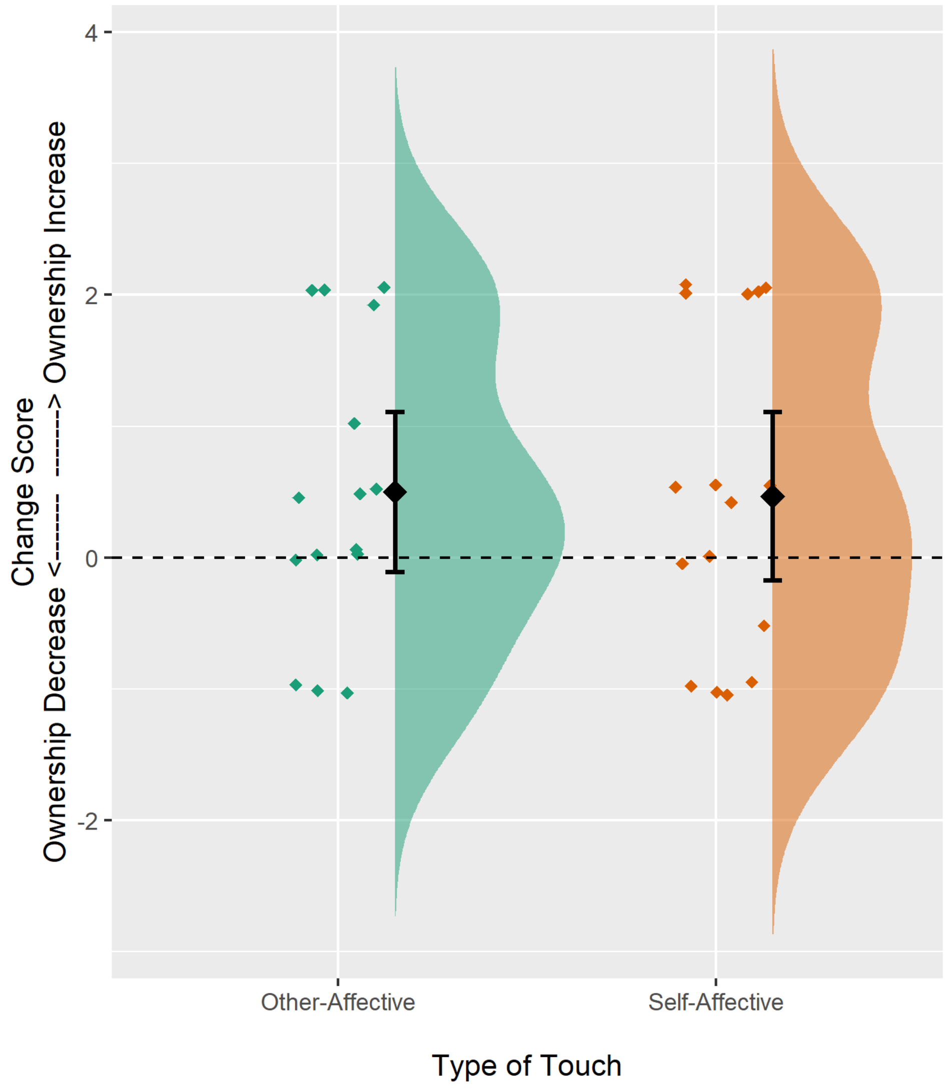

***Figure S2.*** *The effect of touch applied to the right hand on left hand disownership. Positive scores indicate increased ownership. Other-affective touch shown on the left/in green; self-affective touch shown on the right/in orange. Overall distribution of scores represented by the 'cloud'. Raw data indicated by the 'rain' with randomized jitter for improved visualisation. Error bars represent 95% confidence interval around the mean (central diamond).*

This analysis was also run using listwise deletion for missing data and Bayesian statistics, resulting in the same finding that neither type of touch produced a statistically robust change in ownership (listwise deletion Wilcoxon signed-rank test with Bonferroni-corrected $\alpha$=.025: self-affective V(13) = 45.50, p = .280; other-affective V(13) = 37, p = .093. Pairwise deletion Bayesian One-sample T-test: self-affective $BF_{10}$ = .697; other-affective $BF_{10}$ = .907. Listwise deletion Bayesian One-sample T-test: self-affective $BF_{10}$ = .611; other-affective $BF_{10}$ = 1.291). Results of the exact McNemar's test further indicated that a significantly greater proportion of people showed an increase in body ownership following other-affective touch to the left arm compared with the right arm (p = .039).

#### 4.5. Relationship between body ownership (change), touch intensity, and pleasantness ratings

Intensity and pleasantness ratings for different types of touch applied to the left arm are summarised in Supplementary Table S3.

***Table S3.*** *Intensity and Pleasantness ratings by touch condition.*

| | Median | IQR | N |
|---|---|---|---|
| **Intensity (range: 0-10)** | | | |
| Self-Slow | 2.31 | 1.38 – 5.94 | 20 |
| Self-Fast | 3.25 | 1.78 – 6.66 | 20 |
| Other-Slow | 1.5 | 0 – 3.25 | 19 |
| Other-Fast | 1.63 | 0.19 – 4.0 | 20 |
| **Pleasantness (range: 0-5)** | | | |
| Self-Slow | 2.13 | 1.19 – 3.49 | 18 |
| Self-Fast | 2.5 | 1.5 – 3.25 | 17 |
| Other-Slow | 2.0 | 1.50 – 3.0 | 11 |

| | | | |
|---|---|---|---|
| Other-Fast | 2.0 | 1.0 – 3.0 | 15 |

*Note: median and IQR (25th percentile – 75th percentile) reported given non-normal distribution of data.*

Statistical analyses were conducted to identify any differences in the perception of touch intensity and pleasantness across the different types of touch conditions (see Supplementary Figure S3). Inspection of plots indicated that the data were not normally distributed, and so non-parametric (Frequentist) Wilcoxon signed-rank tests were used to examine the main effect of Agent (i.e. by comparing self vs. other touch irrespective of velocity), main effect of velocity (i.e. by comparing slow vs. fast touch irrespective of agent) and interaction between agent and velocity (i.e. by calculating the difference between slow and fast velocity stroking for self touch and separately for other touch and subsequently comparing these differentials). Confirmatory analysis was performed using 2 (Agent: Self vs. Other) x 2 (Touch Velocity: Slow vs. Fast) repeated-measures Bayesian and Frequentist ANOVA, with intensity and pleasantness ratings as dependent variables.

For the *intensity ratings*, Wilcoxon signed-ranks tests identified a main effect of Agent, with self touch rated as significantly more intense than other touch ($W = 511$, $p < .001$), but no main effect of velocity ($W = 171$, $p = .473$) or agent x velocity interaction ($W = 87$, $p = .948$). These results were confirmed by both frequentist ANOVA (main effect of Agent: $F(1,18) = 24.87$, $p < .001$; main effect of Velocity: $F(1, 18) = 1.73$, $p = .21$, ns; Agent x Velocity interaction: $F(1,18) = 0.29$, $p = .60$, ns) and Bayesian analysis (Agency $BF_{10} = 217.76$; Velocity $BF_{10} = .44$; Agency x Velocity interaction $BF_{10} = .34$).

The *pleasantness ratings* showed no significant main effects or interactions using non-parametric or parametric (Frequentist) analyses (all $p > 0.10$), which was substantiated by Bayesian ANOVA, which indicated some support for the null hypothesis of no Agency effect

($BF_{10}$ = .31) or Velocity effect ($BF_{10}$ = .32) (although these values are not below the 0.3 cut-off recommended by Dienes, 2014), and an inconclusive Agency x Velocity interaction ($BF_{10}$ = .57).

Conducting the same analyses removing patients who showed an above-chance tendency to falsely report touch on the left (affected) arm when none was present (i.e. on more than 50% of sham trials; n = 3) did not change any results.

(a)

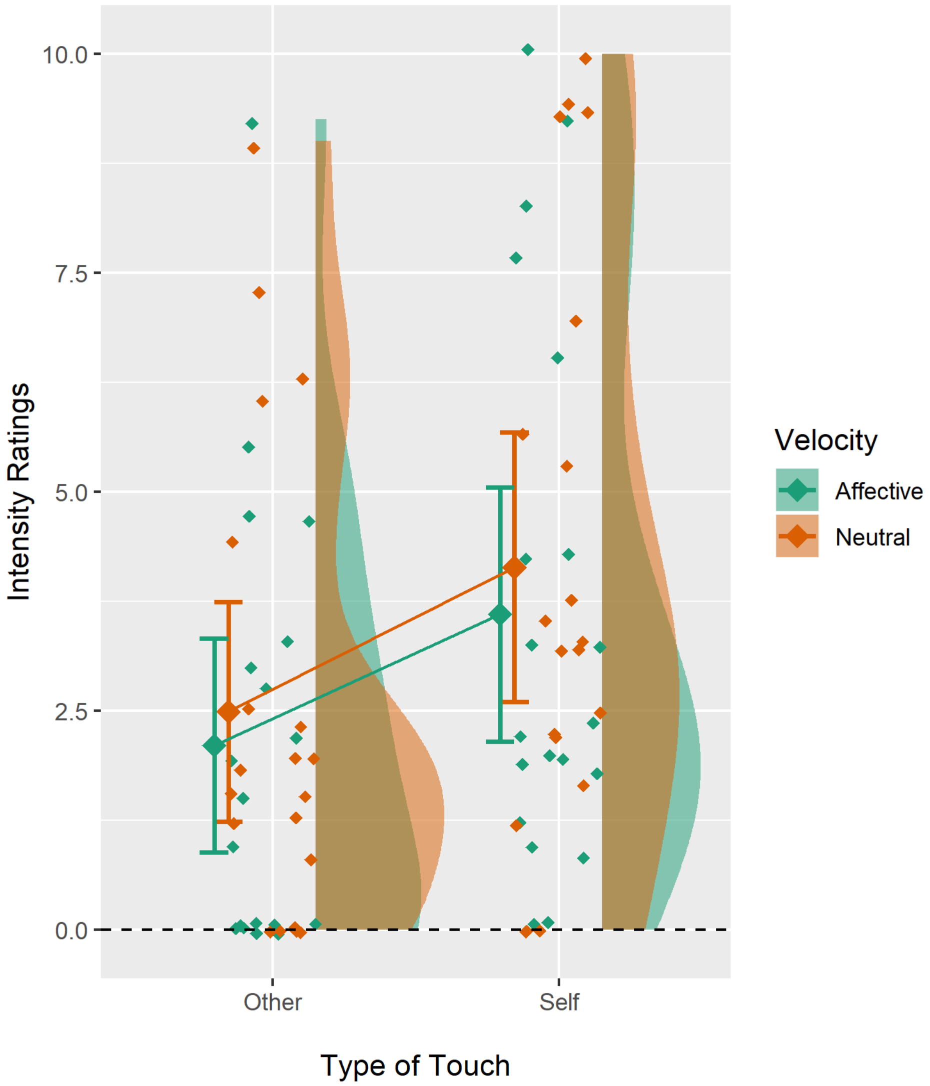

Intensity Ratings
10.0
7.5
5.0
2.5
0.0
Other
Self
Type of Touch
Velocity
Affective
Neutral

(b)

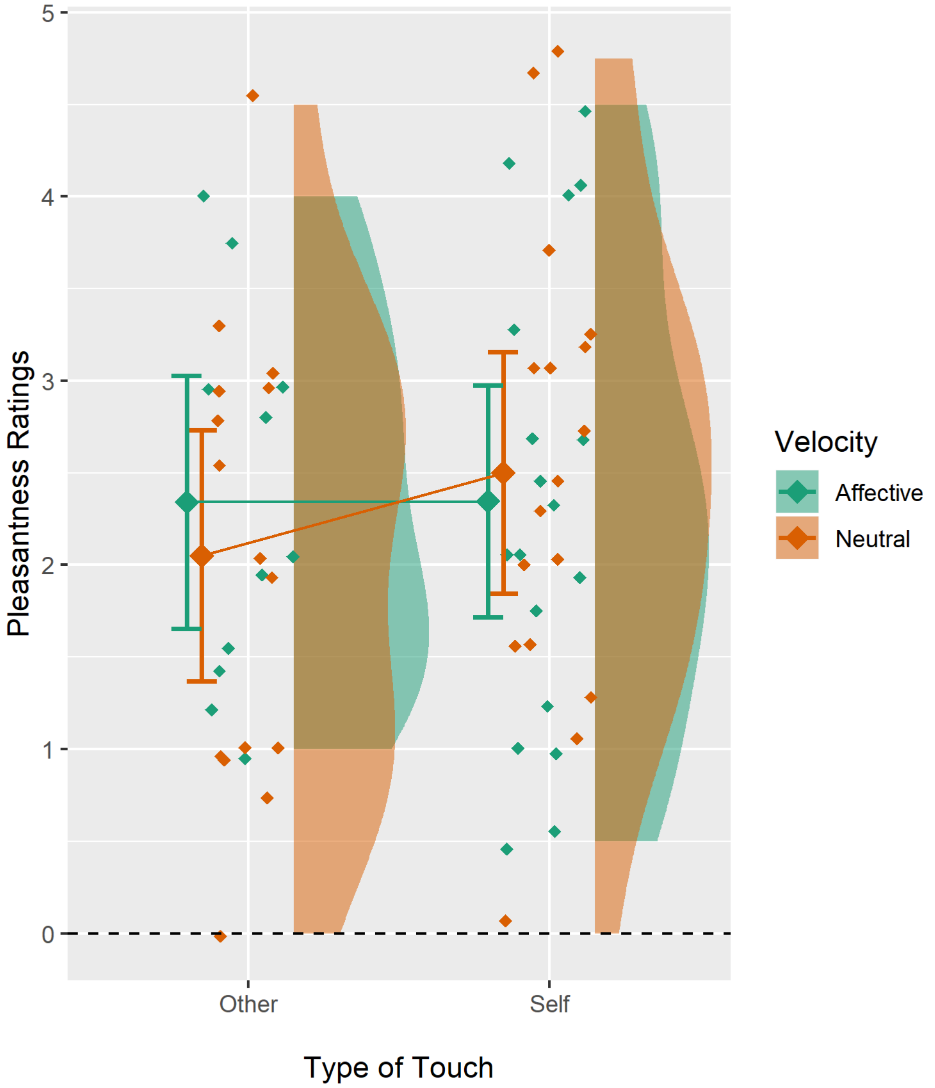

***Figure S3.*** *(a) Intensity ratings (range 0-10), (b) pleasantness ratings (range 0-5). Overall distribution of scores represented by the 'cloud'. Raw data indicated by the 'rain' with randomized jitter for improved visualisation. Error bars represent 95% confidence interval around the mean (central diamond).*

Bivariate (Spearman)'s correlations were used to examine the relationship between baseline DSO scores and touch intensity / pleasantness ratings for each of the four types of touch, as well as between body ownership change scores and the different types of touch.

These did not reveal any significant relationships (all $p > 1.0$). Bayesian correlations similarly did not support any relationship between touch intensity or pleasantness ratings and baseline DSO or ownership change scores (all $BF_{10} < 3.0$). See Supplementary Tables S4-S7 for correlation matrices.

***Table S4.*** *Relationship (Spearman's correlation; $r_s$) between baseline level of DSO and each type of touch in terms of (a) intensity (upper half of table) and (b) pleasantness (lower half).*

| **Rating type** | **Touch type** | **$r_s$, p** |
|---|---|---|
| *Intensity* | Self-Affective | $r_s$ = -0.106, p = 0.655 |
| | Other-Affective | $r_s$ = 0.225, p = 0.355 |
| | Self-Neutral | $r_s$ = 0.246, p = 0.295 |
| | Other-Neutral | $r_s$ = 0.023, p = 0.924 |
| *Pleasantness* | Self-Affective | $r_s$ = -0.027, p = 0.915 |
| | Other-Affective | $r_s$ = -0.376, p = 0.254 |
| | Self-Neutral | $r_s$ = 0.080, p = 0.760 |
| | Other-Neutral | $r_s$ = 0.103, p = 0.716 |

* $p < .05$ uncorrected for multiple tests.

***Table S5.*** *Relationship (Spearman's correlation; $r_s$) between degree of body ownership change and ratings of (a) intensity (upper half of table) and (b) pleasantness (lower half) for each type of touch.*

| | **Ownership Change Following:** | | | |
|---|---|---|---|---|
| | Self-Affective | Other-Affective | Self-Neutral | Other-Neutral |
| **Intensity Rating for:** | | | | |
| Self-Affective | $r_s$ = 0.246, p = 0.295 | $r_s$ = 0.149, p = 0.555 | $r_s$ = -0.147, p = 0.536 | $r_s$ = -0.110, p = 0.646 |
| Other-Affective | $r_s$ = 0.225, p = 0.354 | $r_s$ = 0.138, p = 0.585 | $r_s$ = 0.069, p = 0.780 | $r_s$ = -0.190, p = 0.437 |
| Self-Neutral | $r_s$ = 0.347, p = 0.134 | $r_s$ = 0.112, p = 0.657 | $r_s$ = 0.032, p = 0.893 | $r_s$ = -0.157, p = 0.509 |
| Other-Neutral | $r_s$ = 0.300, p = 0.199 | $r_s$ = 0.136, p = 0.591 | $r_s$ = -0.064, p = 0.787 | $r_s$ = -0.083, p = 0.729 |
| **Pleasantness rating for:** | | | | |
| Self-Affective | $r_s$ =-0.181, p = 0.473 | $r_s$ = -0.041, p = 0.880 | $r_s$ = -0.226, p = 0.367 | $r_s$ = -0.301, p = 0.226 |
| Other-Affective | $r_s$ =-0.434, p = 0.183 | $r_s$ = -0.249, p = 0.487 | $r_s$ = -0.383, p = 0.246 | $r_s$ = -0.446, p = 0.169 |
| Self-Neutral | $r_s$ = 0.054, p = 0.838 | $r_s$ = 0.073, p = 0.795 | $r_s$ = 0.076, p = 0.771 | $r_s$ = -0.020, p = 0.939 |
| Other-Neutral | $r_s$ = 0.075, p = 0.790 | $r_s$ = 0.291, p = 0.335 | $r_s$ = 0.231, p = 0.407 | $r_s$ = 0.010, p = 0.971 |

* $p < .05$ uncorrected for multiple tests.

***Table S6.*** *Bayesian correlations (between baseline level of DSO and each type of touch in terms of (a) intensity (upper half of table) and (b) pleasantness (lower half).*

| **Rating type** | **Touch type** | **r, $BF_{10}$** |
|---|---|---|
| *Intensity* | Self-Affective | r = -0.085, BF = 0.325 |
| | Other-Affective | r = 0.183, BF = 0.516 |
| | Self-Neutral | r = 0.232, BF = 0.750 |
| | Other-Neutral | r = 0.000, BF = 0.286 |
| *Pleasantness* | Self-Affective | r = -0.033, BF = 0.305 |
| | Other-Affective | r = -0.319, BF = 0.868 |
| | Self-Neutral | r = 0.075, BF = 0.335 |
| | Other-Neutral | r = 0.100, BF = 0.370 |

*Note: $BF_{10}$ >3.0 indicates moderate support for the alternative hypothesis. $BF_{10}$ <0.3 indicates moderate support for the null hypothesis. $BF_{10}$ values between 3.0 and 0.3 are inconclusive.*

***Table S7.*** *Relationship (Bayesian correlation) between degree of body ownership change and ratings of (a) intensity (upper half of table) and (b) pleasantness (lower half) for each type of touch.*

| | **Ownership Change Following:** | | | |
|---|---|---|---|---|
| | Self-Affective | Other-Affective | Self-Neutral | Other-Neutral |
| **Intensity Rating for:** | | | | |
| Self-Affective | r = 0.195<br>BF = 0.565 | r = 0.128<br>BF = 0.389 | r = -0.128<br>BF = 0.383 | r = -0.095<br>BF = 0.335 |
| Other-Affective | r = 0.165<br>BF = 0.463 | r = 0.116<br>BF = 0.371 | r = 0.059<br>BF = 0.310 | r = -0.143<br>BF = 0.414 |
| Self-Neutral | r = 0.227<br>BF = 0.720 | r = 0.053<br>BF = 0.314 | r = 0.012<br>BF = 0.286 | r = -0.125<br>BF = 0.379 |
| Other-Neutral | r = 0.224<br>BF = 0.703 | r = 0.094<br>BF = 0.345 | r = -0.056<br>BF = 0.302 | r = -0.073<br>BF = 0.314 |
| **Pleasantness rating for:** | | | | |
| Self-Affective | r = -0.159<br>BF = 0.448 | r = -0.039<br>BF = 0.324 | r = -0.177<br>BF = 0.491 | r = -0.239<br>BF = 0.737 |
| Other-Affective | r = -0.310<br>BF = 0.830 | r = -0.180<br>BF = 0.497 | r = -0.293<br>BF = 0.762 | r = -0.310<br>BF = 0.830 |
| Self-Neutral | r = 0.000<br>BF = 0.308 | r = 0.045<br>BF = 0.335 | r = 0.068<br>BF = 0.330 | r = -0.008<br>BF = 0.309 |
| Other-Neutral | r = 0.067<br>BF = 0.346 | r = 0.216<br>BF = 0.566 | r = 0.181<br>BF = 0.493 | r = 0.045<br>BF = 0.335 |

*Note: $BF_{10}$ >3.0 indicates moderate support for the alternative hypothesis. $BF_{10}$ <0.3 indicates moderate support for the null hypothesis. $BF_{10}$ values between 3.0 and 0.3 are inconclusive.*

### 5. Main Experiment - Lesion Analysis Methods

Clinically acquired CT and MRI scans were used to map lesion location and extent for each patient, following previously established procedures for lesion demarcation (Moro *et al.*, 2016; Martinaud *et al.*, 2017). Briefly, this involved (1) using MRIcron (http://www.nitrc.org/projects/mricron) to rotate a high-resolution, MRI template (in terms of yaw, pitch and roll) to match the orientation of the patient's native scan, after which (2) the extent of each lesion (including visible oedema) was manually draw on to the template by an experienced researcher, using established landmarks as a guide, and checked by a second, senior researcher (VM) before, (3) the template and lesion map was re-oriented back to standard space ready for further analysis. We chose to follow this established, manualised method rather than alternative, automated lesion demarcation procedures, owning to current limitations of automated procedures (see Maier *et al.*, 2015; de Haan and Karnath, 2018; Liew *et al.*, 2018), which are especially apparent when using CT scans.

Subsequently, a lesion overlay of all patients was created, before univariate voxel-wise lesion-symptom mapping (VLSM) was implemented using NPM (non-parametric mapping; http://www.cabiatl.com/mricro/npm/) to identify anatomical regions associated with (1) disownership scores obtained during the baseline (pre-touch) condition of the experimental task (n = 24; scores inverted to adhere to the requirement of data input directionality, such that higher scores indicate better performance [i.e. greater body ownership]), and (2) ownership change scores obtained from the other-affective touch condition (n = 18 patients with baseline disownership >0 and available behavioural plus lesion data). We used the Brunner-Menzel test for non-normally distributed behavioural data and permutation thresholding ($p < 0.05$ using 4000 permutations) to correct for multiple comparisons. To ensure sufficient statistical power and reduce spatial bias, we excluded voxels damaged in fewer than 10% of the patient sample, and in preliminary analysis we examined the relationship between lesion volume and the behavioural scores (de Haan and Karnath, 2018; Sperber and Karnath, 2018), finding no such relationship (see results). In subsequent, less restrictive analyses (stemming from non-significant results using the methods above), we performed the same VLSM including all voxels (not only those lesioned in at least 10% of patients), and a less restrictive correction criteria of 1% False Discovery Rate.

The resulting statistical lesion maps were superimposed onto grey and white matter anatomical atlases (i.e. the automated anatomical labelling [AAL] template a high-resolution template, Tzourio-Mazoyer *et al.*, 2002; the NatBrainLab template of human brain connections, Catani *et al.*, 2013) using MRIcron, for visualisation and anatomical interpretation.

### 6. Main Experiment - Lesion Analysis Results

Preliminary analyses showed that lesion volume did not correlate with baseline DSO scores ($r_s$ (24) = .137, p = .523), nor other-affective touch ownership change scores ($r_s$ (18) = -0.199, p = .428), and so was not considered a nuisance variable in subsequent analyses. The VLSM of lesions associated with baseline experimental body disownership scores did not show any significant results in our sample using the stringent parameters detailed above (BM z-range = -3.13 to 2.12; permutation-corrected $p < .05$ for $z > 3.72$), nor the less restrictive criteria (no minimum lesion overlap, and 1% FDR correction; $p < 0.01$ for $z > 9.20$).

The VLSM of lesions associated with body ownership change following affective touch likewise did not survive permutation based statistical thresholding (BM z-range = -2.25 to 2.91, permutation-corrected $p < .05$ for $z > 3.43$); however, the less conservative, exploratory analysis using no minimum lesion overlap and 1% FDR-corrected threshold identified damage in the right insula cortically, and more substantially the white matter comprising right corpus callosum, and to a smaller extent, cingulum and cortico-spinal tract ($p < .01$ for $z > 2.42$; see Supplementary Table S8). Damage to these areas was associated with a failure for limb ownership to increase following other affective touch. Statistical lesion maps are illustrated in Figure 3 (main document).

***Table S8.*** *Significant voxels from the VLSM where increases in ownership following other-affective touch is significantly lower in patients with damage than ownership increases in patients without damage. The number of voxels for each region indicated in the brain atlas of grey (AAL) and white matter (NatBrainLab) are reported, along with the centre of mass (MNI co-ordinates).*

| **Atlas** | **Area** | **N > 0** | **x, y, z** |
|---|---|---|---|
| AAL | Insula | 16 | 35, 4, 13 |
| Nat BrainLab | Corpus callosum | 61 | 4, -3, 24 |
| | Cingulum | 2 | 8, 0, 26 |
| | Cortico spinal tract | 1 | 20, -7, 2 |

**Supplementary Materials References**

Bisiach E, Vallar G, Perani D, Papagno C, Berti A. Unawareness of disease following lesions of the right hemisphere: Anosognosia for hemiplegia and anosognosia for hemianopia. Neuropsychologia 1986; 24: 471–482.

Catani M, Thiebaut de Schotten M, Catani M, Thiebaut de Schotten M. Atlas of Human Brain Connections (all tracts). In: Atlas of Human Brain Connections. 2013

Gentsch A, Panagiotopoulou E, Fotopoulou A. Active Interpersonal Touch Gives Rise to the Social Softness Illusion. Curr Biol 2015; 25: 2392–2397.

de Haan B, Karnath HO. A hitchhiker's guide to lesion-behaviour mapping. Neuropsychologia 2018

Halligan PW, Cockburn J, Wilson BA. The behavioural assessment of visual neglect. Neuropsychol Rehabil 1991; 1: 5–32.

Kirsch LP, Besharati S, Papadaki C, Crucianelli L, Bertagnoli S, Ward N, et al. Damage to the Right Insula Disrupts the Perception of Affective Touch. bioRxiv 2019: 592014.

Liew SL, Anglin JM, Banks NW, Sondag M, Ito KL, Kim H, et al. A large, open source dataset of stroke anatomical brain images and manual lesion segmentations. Sci Data 2018

Maier O, Schröder C, Forkert ND, Martinetz T, Handels H. Classifiers for ischemic stroke lesion segmentation: A comparison study. PLoS One 2015

Martinaud O, Besharati S, Jenkinson PM, Fotopoulou A. Ownership illusions in patients with body delusions: Different neural profiles of visual capture and disownership. Cortex 2017; 87

von Mohr M, Kirsch LP, Loh JK, Fotopoulou A. Affective Touch Dimensions: From Sensitivity to Metacognition. bioRxiv 2019: 669259.

Moro V, Pernigo S, Tsakiris M, Avesani R, Edelstyn NMJ, Jenkinson PM, et al. Motor versus body awareness: Voxel-based lesion analysis in anosognosia for hemiplegia and somatoparaphrenia following right hemisphere stroke. Cortex 2016; 83: 62–77.

Sperber C, Karnath HO. On the validity of lesion-behaviour mapping methods. Neuropsychologia 2018

Tzourio-Mazoyer N, Landeau B, Papathanassiou D, Crivello F, Etard O, Delcroix N, et al. Automated Anatomical Labeling of Activations in SPM Using a Macroscopic Anatomical Parcellation of the MNI MRI Single-Subject Brain. Neuroimage 2002; 15: 273–289.
